# Establishment of an optimized and automated workflow for whole brain probing of neuronal activity

**DOI:** 10.1101/2024.09.16.611953

**Authors:** Sébastien Cabrera, Nicolas Vachoud, Renato Maciel, Marine Breuilly, Stevenson Desmercieres, Geraldine Meyer-Dilhet, Salma Ellouze, Julien Courchet, Pierre-Hervé Luppi, Nathalie Mandairon, Olivier Raineteau

## Abstract

Behaviors are encoded by widespread neural circuits within the brain that change with age and experience. Immunodetection of the immediate early gene c-Fos has been successfully used for decades to reveal neural circuits active during specific tasks or conditions. Our objectives here were to develop and benchmark a workflow that circumvents classical temporal and spatial limitations associated with c-Fos quantification. We combined c-Fos immunohistochemistry with c-Fos driven Cre-dependent tdTomato expression in the TRAP2 mice, to visualize and perform a direct comparison of neural circuits activated at different times or during different tasks. By using open-source software (QuPath and ABBA), we established a workflow that optimize and automate cell detection, cell classification (e.g. c-Fos *vs*. c-Fos/tdTomato) and whole brain registration. We demonstrate that this workflow, based on fully automatic scripts, allows accurate cell number quantification with minimal interindividual variability. Further, interrogation of brain atlases at different scales (from simplified to detailed) allows gradually zooming on brain regions to explore spatial distribution of activated cells. We illustrate the potential of this approach by comparing patterns of neuronal activation in various contexts (two vigilance states, complex behavioral tasks…), in separate groups of mice or at two time points in the same animals. Finally, we explore software (BrainRender) for intuitive representation of the results. Altogether, this automated workflow accessible to all labs with some expertise in histology, allows an unbiased, fast and accurate analysis of the whole brain activity pattern at the cellular level, in various contexts.

## INTRODUCTION

For decades, researchers have probed the distribution of neuronal activity that drives specific behaviors, aiming to unravel the complexity of brain functional organization. To this end, the detection of immediate-early genes (IEGs) has been widely used to spatially monitor behaviorally-induced neuronal activity^1,2^. In recent years, the neuroscientific community has witnessed the emergence of whole brain quantifications and analyses, driven by the emergence of brain clearing and whole brain staining procedures^3–5^. However, these approaches require a high level of expertise in tissues processing, image acquisitions, and analysis, which are not accessible to most research teams. Alternative approaches are based on immunohistochemistry, classical stereological rules (homogeneous brain sampling) and automatic image acquisition (i.e. slide scanner) which are routinely used in many laboratories. Nevertheless, beyond the initial steps of section processing and image acquisition, two critical phases necessitate optimization for robust analysis: 1) cellular segmentation, which involves delineating individual cells based on single or multiple markers’ expression, and 2) section registration onto a reference atlas. To address these challenges, dedicated software have emerged to optimize both segmentation and registration processes, enabling efficient analysis across a large number of sections ^6,7^. These software enhance the accessibility of these sophisticated analyses and also facilitate their reproducibility and scalability.

Here, we established a workflow for performing whole brain estimates of cell counting, exclusively using open source software (QuPath/ImageJ and ABBA). We optimized and benchmarked all essential steps necessary for accurate cell quantification, enabling robust comparisons between experimental conditions. we provide scripts for user-friendly implementation of the workflow, incorporating quality control assessments at critical steps. We illustrate the efficacy of this workflow by using immunodetection of c-Fos, a widely recognized marker of brain activity, alongside a recent transgenic mouse model (i.e. *TRAP mouse*) which allows conditional expression of a reporter protein in c-Fos expressing cells. The integration of these tools enables the study of brain activity associated with distinct behaviors within the same animal, thus representing a refinement in adherence to the 3Rs principles of animal research.

Altogether, our work provides a comprehensive workflow for whole-brain mapping of neuronal activity based on IEGs expression. This workflow can be executed without the need for specialized equipment, facilitating the generation of structure-function hypotheses that can be further tested by using complementary approaches.

## RESULTS

### Development of the protocol

This protocol establishes a workflow for cellular segmentation based on unique or multiple markers, followed by section registration and analysis, including 3D visualization and statistical interpretation of acquired data. It is designed for experimental biologists interested in analyzing and comparing patterns of activity, based on c-Fos immunodetection or tdTomato detection in the TRAP2 mouse, across diverse behavioral contexts. It relies on open source software, as well as on histological techniques that are mastered by most research labs, thereby ensuring accessibility and reproducibility across research laboratories. All steps of the workflow were benchmarked to define optimal parameters for cell detection and atlas registration. Fully annotated scripts are provided, allowing quality control at critical steps of the workflow, to maintain high standards throughout the analyses. We analyzed c-Fos expression, as a marker of neuronal activity, however, our workflow is applicable to any nuclear markers and can also be applied to cytoplasmic markers following minor adjustments, broadening its applicability across diverse research domains.

### Overview of the protocol

This section outlines the major steps of the workflow (see **Figure 1**).

**Fig 1:**
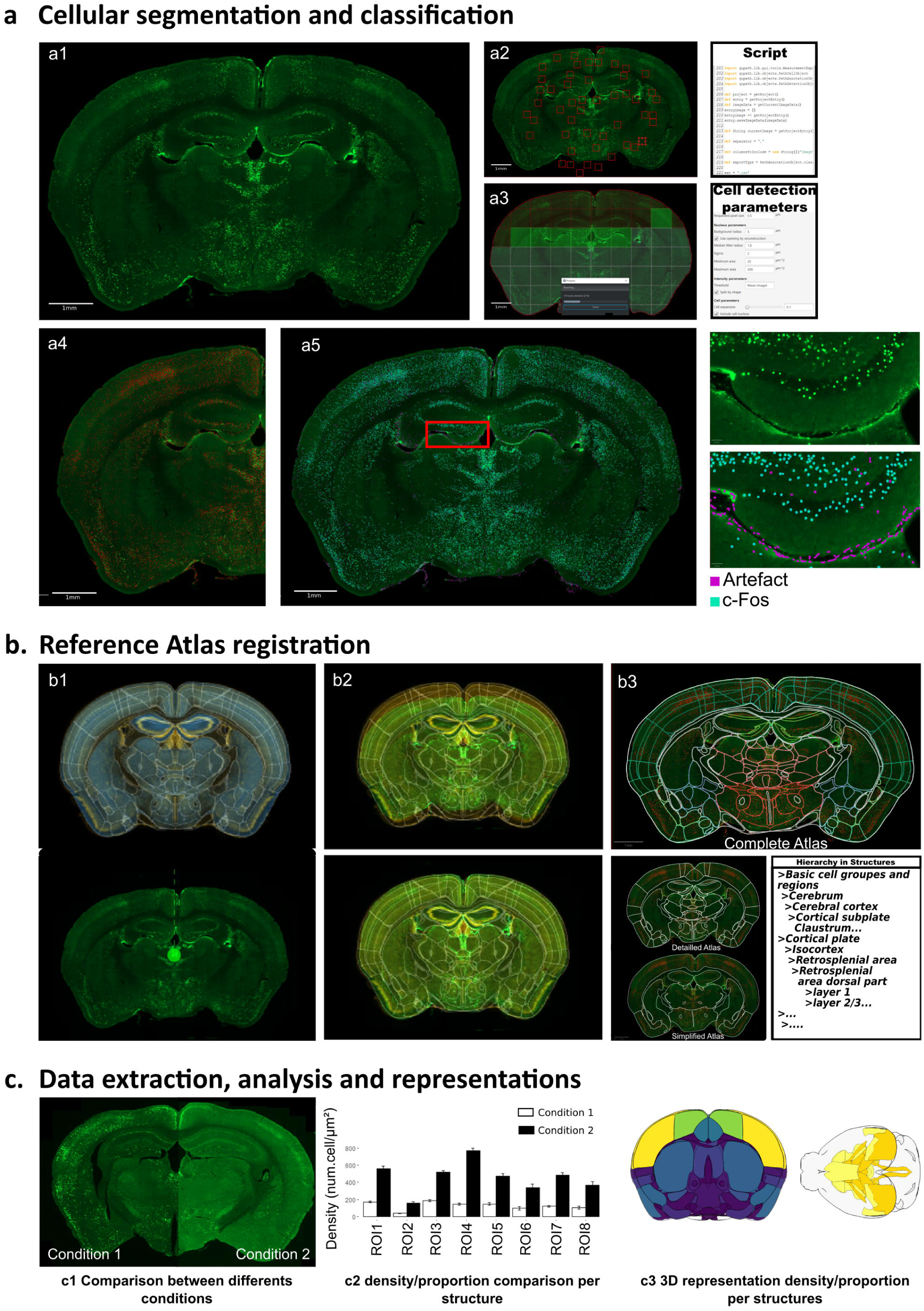
Representation of the complete workflow for cell segmentation and classification, atlas registration and analysis of cell distribution. The workfiow is composed of three consecutive steps: **a**, cellular segmentation and classification using an automated script in QuPath; **b**, reference atlas registration with ABBA software; and **c**, data extraction, analysis and representation. **a)** Cellular segmentation begins with c-Fos immunostaining on serial sections, which are then imaged on a slide scanner **(a1)**. Adaptive thresholding is performed using a script that extracts values from random fields **(a2)**, then applies it to the entire section **(a3)** for optimal cell detection.Once cell detection is complete **(a4)**, a classifier is used **(a5)** to differentiate c-Fos positive cells (blue) from artifacts (purple), as illustrated in a higher magnification of the dentate gyrus. **b)** During the second step, ABBA software is used to distribute and register individual sections onto the Allen reference brain atlas **(b1)**. Elastic algorithms are used to improve section adaptation to the reference atlas, followed by final manual refinement **(b2)**. Different atlas resolution levels can be generated based on the hierarchical organization of the Allen mouse brain atlas **(b3)**. **c)** Data analysis begins with quality control for individual mice, followed by intra- and inter-groups comparisons using a R script provided as supplementary material **(c1)**. Different

<u>Stage 1 - Cellular segmentation and classification (</u>**Figure 1a**): it relies on QuPath^8^ (https://qupath.github.io/). Scripts for automation and adaptive thresholding using Qupath or ImageJ are provided. Both result in accurate detection of labeled cells while reducing variability observed between sections. Classifiers to detect and subtract artifacts were also benchmarked and included into the workflow.

<u>Stage 2 - Reference atlas registration using ABBA (</u>**Figure 1b**): it relies on ABBA (Aligning Big Brain Atlases, https://abba-documentation.readthedocs.io/). Essential principles for optimization of the registration procedure are provided.

<u>Stage 3 - Data extraction, analysis and representation (</u>**Figure 1c**): analysis is performed in R in two steps (i.e., 1) quality control of individual animals and 2) comparison of animal groups corresponding to two independent scripts. A third script allows representing results in a 2D/3D model by using Brain Render (https://github.com/brainglobe/brainrender, ^9^).

### Histology & Imaging

We aimed at developing a universally applicable methodology for comparing results produced by different laboratories or produced under varying experimental conditions. To this end, we used two sections thickness (30 and 50 micrometers), encompassing sections generated via cryostat or vibratome methods. Notably, we observed no discernible differences in section quality or immunostaining efficacy between the two techniques. In both cases, staining was performed on free-floating sections. Thus, the workflow described below can be applied to a range of sectioning/staining procedures commonly used in diverse laboratory settings. However, for optimal whole brain analysis, several rules should be strictly followed. For whole brain estimates, it is absolutely essential to follow strict stereological rules for tissue collection. Indeed, homogeneous tissue sampling is important for unbiased quantifications (https://www.stereology.info/sampling/). Given the considerable variation in volume across brain structures, we recommend a section interval of 250µm or less for c-Fos quantification, owing to the abundance of c-Fos positive cells. For particularly small brain structures or for antigens exhibiting sparse labeling, a higher sampling rate (i.e. smaller section interval), may be necessary. In our analysis, we chose to collect 1 section every 8 sections in paradoxical sleep (PS) and wakefulness (WF) conditions (thickness: 30µm; section interval: 210µm) and 1 section every 6 in social behavior (SB) condition (thickness: 50µm; section interval: 250µm).

For imaging, we recommend using a slide scanner (here an Axioscan, Zeiss). Optimal acquisition parameters tailored to the specific experimental conditions, have to be determined by the user. Settings should be balanced to avoid saturated or weak signal detection. For thick sections such as those used in this work, multiple focal planes should be imaged to ensure full visualization. Depending on staining of interest, specific functions available in Zen software (Zeiss) can be applied to improve scan sharpness, thereby facilitating quantification precision (e.g. using the wavelet function for c-Fos/tdtomato staining). Finally, we recommend exporting single section mosaics using the “processing-batch-split-scene” function (Zen software, Zeiss) at the end of the acquisition, rather than loading the full .czi file containing multiple sections in Qupath. This practice is particularly useful when dealing with sections from multiple wells (e.g., wells 1 and 7 of a series of 12, for obtaining a 1:6 sampling rate), allowing for the orderly arrangement of sections before importing them into Qupath. Indeed, correct ordering of sections is required to facilitate registration (see stage 2) and subsequent analysis (see stage 3).

### Stage 1: Cellular segmentation and classification using QuPath

We first aimed at assessing the accuracy of c-Fos detection using QuPath (**Figure 2a**). To this end, we determined the range of c-Fos immunoreactivity intensity across randomly selected sections and tested three levels of threshold (low, 420; medium, 3274; high, 6127) allowing us to detect all cells or to restrict the detection to those more strongly stained. Visual inspection confirmed the precision of the detection process, with as expected, an inverse correlation between threshold levels and number of cells detected (**Figure 2b**). Accordingly, the average c-Fos intensity was of 9328 ± 356 gray values using the “low threshold”, resulting in around 24067 ± 1022 c-Fos positive cells being detected per section. This average gray value increased with “medium threshold” to 18452 ± 585 and “high threshold” to 24144 ± 672, resulting in fewer cells being detected (respectively 8308 ± 442 cells and 4746 ± 349). Thus, QuPath allows accurate detection of c-Fos expressing cells while providing flexibility to adjust the detection sensitivity to detect cells with different staining intensities.

**Fig 2:**
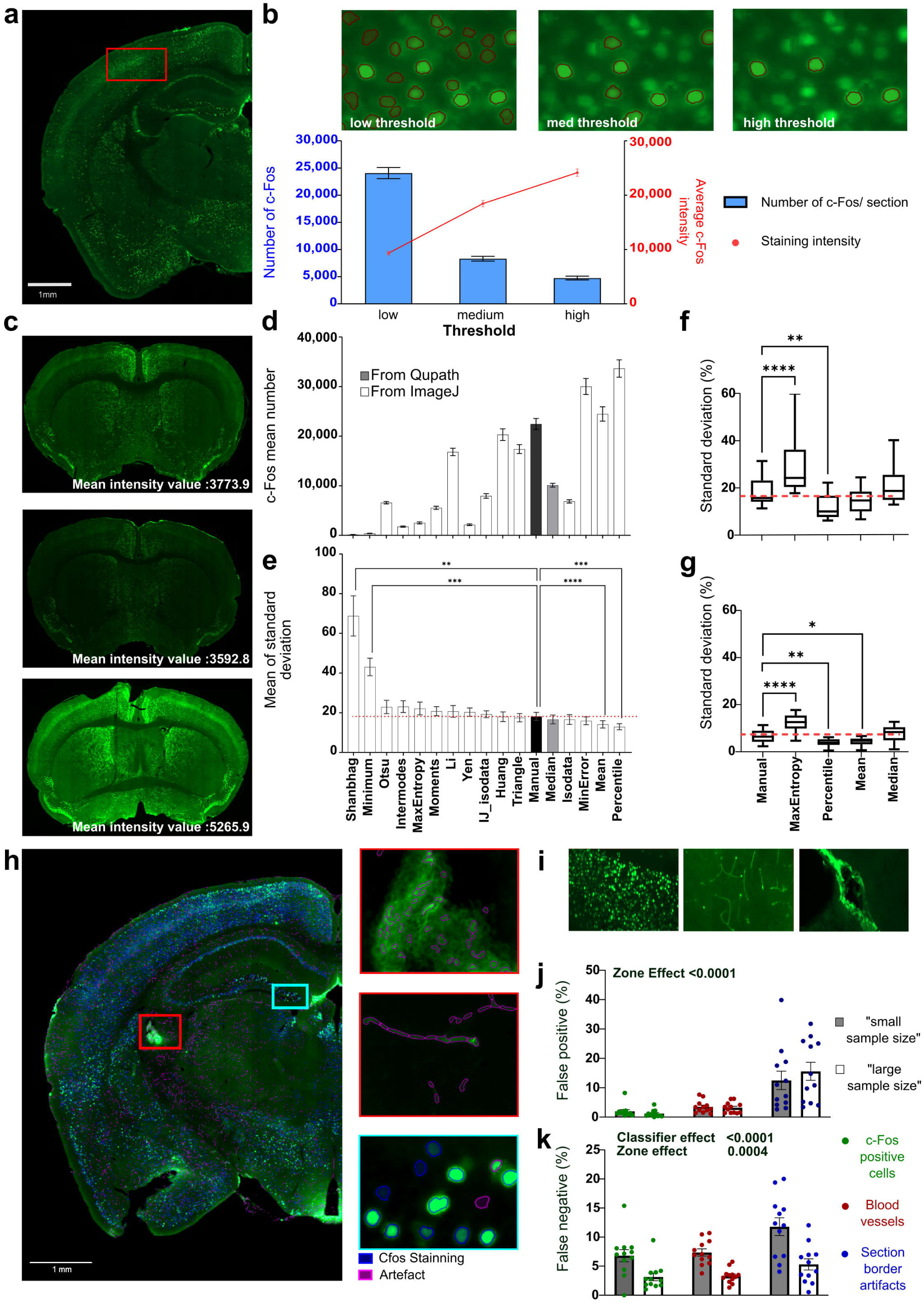
Automation and refinement of c-Fos positive cell detection and classification. **a)** Representative section immunostained for c-Fos. Captions illustrate how detection at 3 arbitrary threshold levels, i.e. low (420), medium (3274) and high (6127), impacts the number of detected cells. Data represent n=6 brain sections per threshold level. **b)** Quantification of detected cell number and averaged intensities at the three arbitrary thresholds (grey level). **c)** Illustration of pronounced staining intensity variability between subsequent sections. **d)** Quantification of c-Fos positive cells using manual arbitrary thresholding (white bar, 1000) or various automatic thresholding methods in QuPath (grey bar) or lmageJ (line-patterned bars). Note the variability in detection efficiency. Data represent n=14 brain sections per thresholding method. **e)** Quantification of count variability (i.e. standard deviation) between three consecutive sections using manual arbitrary thresholding (white bar) or various automatic thresholding methods in QuPath (blue bar) or lmageJ (pink bars). Data represent n=14 brain sections per thresholding method. **f-g)** Selection of automatic thresholding methods based on detection efficiency and reduced count variability (shown in d and e) for complementary analysis of performance with consecutive sections showing heterogeneous **(f)** and homogeneous **(g)** staining. Standard deviation of counts obtained from section triplicates are shown. The “Percentile”, “mean” and “median” methods can be used for automatic thresholding, offering various detection sensitivities and similar or reduced count variability compared to manual arbitrary thresholding (red line, 1000). Data represents n=38 sections per thresholding method. **h)** Representative section showing different sources of artifacts (purple), i.e. blood vessels and border effects. **i-k).** Comparison of two “training” methods for distinguishing artifacts from positive cells using classifiers (i.e. “large sample size” and “small sample size”, see methods). Manual validation of classification errors in identifying positive cells (j) or artifacts **(k)** in regions with a large number of cells or artifacts (i). Data represent n=12 samples per classifier analyzed and region analyzed. Note that both methods show comparable efficiency in accurately detecting positive cells, but the “large sample size” training method significantly reduces errors while detecting

Quantitative analysis of c-Fos immunodetection faces two primary challenges that impact accuracy: 1) variability of staining intensity within section, and 2) variability of staining intensity across sections induced by the immunostaining method used (floating sections, *see material and methods*). Although manual adjustment of detection thresholds, as demonstrated earlier, can mitigate these challenges, unbiased methods are preferable. While the first source of variability may carry biological significance, the second is largely experiment-dependent, resulting for example from tissue folding or adherence to the vessel walls during staining. QuPath only offers limited capabilities for adaptive thresholding, but mostly relies on experimentator-defined values. We therefore explored different means to perform automatic thresholding (**Figure 2c-e**). We evaluated and compared the following approaches: 1) <u>Internal</u> <u>automatic thresholding (QP):</u> this method relies on automatic detection of the median intensity measured within individual sections. A script (“Automatic_threshold_application_(Qupath)”) facilitates the extraction of this variable to define a threshold for cell detection in each section. 2) <u>External automatic thresholding (IJ):</u> this approach relies on thresholding methods implemented in ImageJ. A script (“Automatic_threshold_application_(ImageJ)”) allows defining random fields to define an average staining intensity per section. Notably, performing this detection multiple times considerably improves the accuracy of intensity detection and, consequently, the defined threshold.

To determine the best automatic thresholding methods, we compared results obtained on adjacent sections showing variable staining intensities, using an arbitrary threshold of 1000 (22073 ± 1414 detections per section), to those obtained through 16 automatic thresholding methods (**Figure 2d-e**). Both the mean number of detections per section and the SD between adjacent sections (triplicates) were measured. Results highlight that thresholding methods result in varying levels of detection (**Figure 2d**), ranging from a 48,88% increase (Percentile, 32863 ± 2045 detections) to a 99% decrease (Shangbag, 135 ± 27 detections) compared to manual thresholding. Several methods yielded detection counts comparable to those obtained with the arbitrary threshold, such as the “Huang” method (19931 ± 1371 detections). Among all tested methods, the “Mean” (23988 ± 1641 detections), “Median” (10087 ± 600 detections) and “MaxEntropy” (2531 ± 239 detections) were of interest, as these approaches enabled detecting a range of c-Fos intensities close to the low, medium and high thresholds tested above. We next checked how different threshold methods influenced the variability in cell counts between three consecutive sections, taking as a reference the manual thresholding (**Figure 2e**). Statistical analysis on the standard deviation revealed a significant influence of the thresholding methods on cell counts (F(2,336;30.37)=17.90; p<0.0001; Greenhouse–Geisser ε = 0.1460). We observed that specific automatic threshold methods resulted in increased variability in cell counts compared to the manual thresholding, notably “Shanbhag” (**+300%**, p<0.01, Dunnett’s multiple comparison test). In contrast, “Mean” and “Percentile” decreased this variability (p<0.01, **-20%** for Mean and **-30%** for Percentile) while “Max Entropy’’ and “Median” demonstrated no significant effect. To complement these findings, we selected 4 automatic thresholding methods (“MaxEntropy”, “Percentile”, “Mean” and “Median”) and evaluated in more detail the percentage of variability of cell detection on three consecutive sections showing heterogeneous **(Figure 2f)** and homogenous (**Figure 2g**) cell counts when using a manual threshold of 1000. In heterogeneous sections (**Figure 2f**), we found a similar percentage of standard deviation (SD) for “Mean” (14.39 ± 0.79%, p=0.16) and “Median” methods (21.26 ± 1.24%, p=0.42) compared to the manual thresholding (18.09 ± 0.94%). However, we found a lower percentage of SD with the “Percentile” method (12.32 ± 0.84, p<0.01, −31%) and higher with the “MaxEntropy’’ method (28.02 ± 1.577, p<0.01). In homogeneous sections (**Figure 2g**), we found a similar standard deviation for “Median” methods (7.90 ± 0.50%, p=0.56) and manual threshold (6.64 ± 0.45). The “Percentile” (3.84 ± 0.27%, p<0.01) and “Mean” (4.27 ± 0.27%, p<0.05) methods reduced standard deviation values, while “MaxEntropy’’ increased it (12.19 ± 0.56%, P<0.01, +50% compared to manual threshold). However, despite “percentile” yielding satisfactory outcomes in terms of the number of identified cells and standard deviation, it was discarded due to visual inspection revealing over segmentation, with many artifacts taken into account. Altogether, these results led to the selection of the “mean” (MeanIJ), “median” (MedQP) and “MaxEntropy” (MaxE) methods, based on their capacity to reveal varying levels of detection without exacerbating or even reducing the variability of cell counts.

Cell counting often results in false positive detection, which can arise from blood vessels auto-fluorescence or antibodies aggregation at section borders (**Figure 2h-i**). To address this issue, classifiers using machine learning approaches integrated within QuPath (called *“cell classifier”*) can be used to detect artifactual detection. These classifiers discern between genuine signal and artifacts by manually assigning detected cells to “positive” or “artifact” classes. The efficiency of a classifier relies on training conducted by the experimenter, making it challenging to determine the optimal level. Thus, while undertraining may not be sufficient for optimal classification of artificial signals, overtraining may also result in ambiguous results. We assessed two strategies for training classifiers. The first strategy, termed “Small sample size” involves sampling small areas with few detected cells (<10), repeated numerous times (>90 repeats per detected cell class). Conversely, the second strategy, termed “large sample size”, consists in sampling larger areas with a higher number of detected cells (>50), but fewer repetitions (<10 repeats per detected cells class). To compare both strategies, we manually curated results obtained using each approach to quantify the number of falsely classified cells as “positive” or “artifact” across three distinct areas containing c-Fos positive cells, blood vessels and section border artifacts (n=12 sections per area and classifier across four mice). Results indicate that no changes were noted in the proportion of cells falsely classified as positive (**Figure 2j**, F(1,66) = 0.1941, p=0.6610). However, the “large sample size” strategy resulted in a significant reduction of cells falsely classified as artifact (**Figure 2k**, F(1,66)=38.32, p<0.0001). Altogether, our results underscore the superiority of the “large sample size” strategy, which was therefore used in subsequent analyses.

### Stage 2- Reference atlas registration using ABBA

Several atlases are available online. With the open-access Allen Brain Atlas quickly establishing itself as a reference to most neuroanatomists (http://atlas.brain-map.org, ^10^). Following cell detection and classification, sections can be aligned to this reference atlas using ABBA (Aligning Big Brain Atlases). The registration workflow implemented in this software is detailed online (https://abba-documentation.readthedocs.io/). It facilitates the adaptation of reference brain atlas contours to sections through a combination of automatic (i.e. elastik affine + spline) and manual algorithms (BigWarp^11^) ensuring that deformations are applied solely to the contours, leaving the brain section image and detected cell geometry unaltered. The registration workflow consisted in several steps, which we evaluated to establish the most efficient approach. The initial step involves identifying key sections for optimal rostro-caudal positioning and cutting plane correction. At the beginning of the alignment process, we recommend selecting the most rostral and the most caudal sections. Initial rostro-caudal registration can be achieved by using the maximal section interval offered by the reference atlas. Subsequently, the section interval can be reduced to fine-tune section placement. If sampling is uniform, sections can then be evenly distributed between the two key sections. Validation is then performed by visual inspection of all sections to rectify any potential errors that may have occurred during sectioning (i.e. missing or duplicated sections). ABBA registration further facilitates correction for section inclination along the “x” and “y” axes. Indeed, even if brain sectioning protocols may be standardized using brain casts ^12^, variations across brains are common. For precise cutting plane correction, we recommend the following workflow and landmarks. The “x” axis correction is done by focusing on a caudal section corresponding to section 79-80 of the Allen adult mouse atlas (**Extended data figure 1a**). This section, with a well-defined dorsal hippocampus and the separation of the corpus callosum into two hemispheric structures, is prone to notable “x” axis changes. Positive inclination accelerates the rostro-caudal separation of the corpus callosum, while it persists as a continuous structure on a larger number of sections with negative inclination. The dorsal hippocampus within this section can be used for optimal “y” axis correction, as even a slight inclination in the “y” axis results in well visible differences between the two hippocampi, particularly evident for the dentate gyri (**Extended Data Figure 1a**). Validation of “y” axis correction can then be conducted by selecting a more rostral section (section 53 of the Allen adult mouse atlas, **Extended Data Figure 1b**), where the anterior commissure is continuous between hemispheres, thereby confirming “y” axis correction.

Following establishment of initial parameters, it is recommended to sequentially employ both linear (affine) and nonlinear (spline) “elastik algorithms’’ integrated into the ABBA software, to further refine section registration. This process involves selecting a specific channel of the image (e.g. Dapi, c-Fos… **Figure 3a**) to align it with one of the reference atlases available in ABBA (i.e. Nissl or 3D model “Ara”, **Figure 3b**). To systematically evaluate the efficacy of registration across all channels and reference atlases, we adopted the following approach. We first confirmed improvement in registration accuracy through the application of affine (applied once or twice) and spline transformations, using the c-fos channel and Nissl Atlas. We then compared results obtained using distinct staining channels (including counterstaining for neurons, i.e. NeuN and myelin, i.e. MBP). We finally compared results obtained using the most effective channels with both the Nissl and “Ara” Atlas models. For quantitative assessment, user drawn contours of selected regions were systematically compared against the results achieved by the registration process (n= 8 regions of interest analyzed per combination of parameters evaluated). This comparison allows for the calculation of a Dice coefficient (see script “Dice_contour_calculation” for dice coefficient calculation in QuPath), whose value increases along with contours superposition. A Dice coefficient greater than 0.7 indicates acceptable contours superposition^13^.

**Fig 3:**
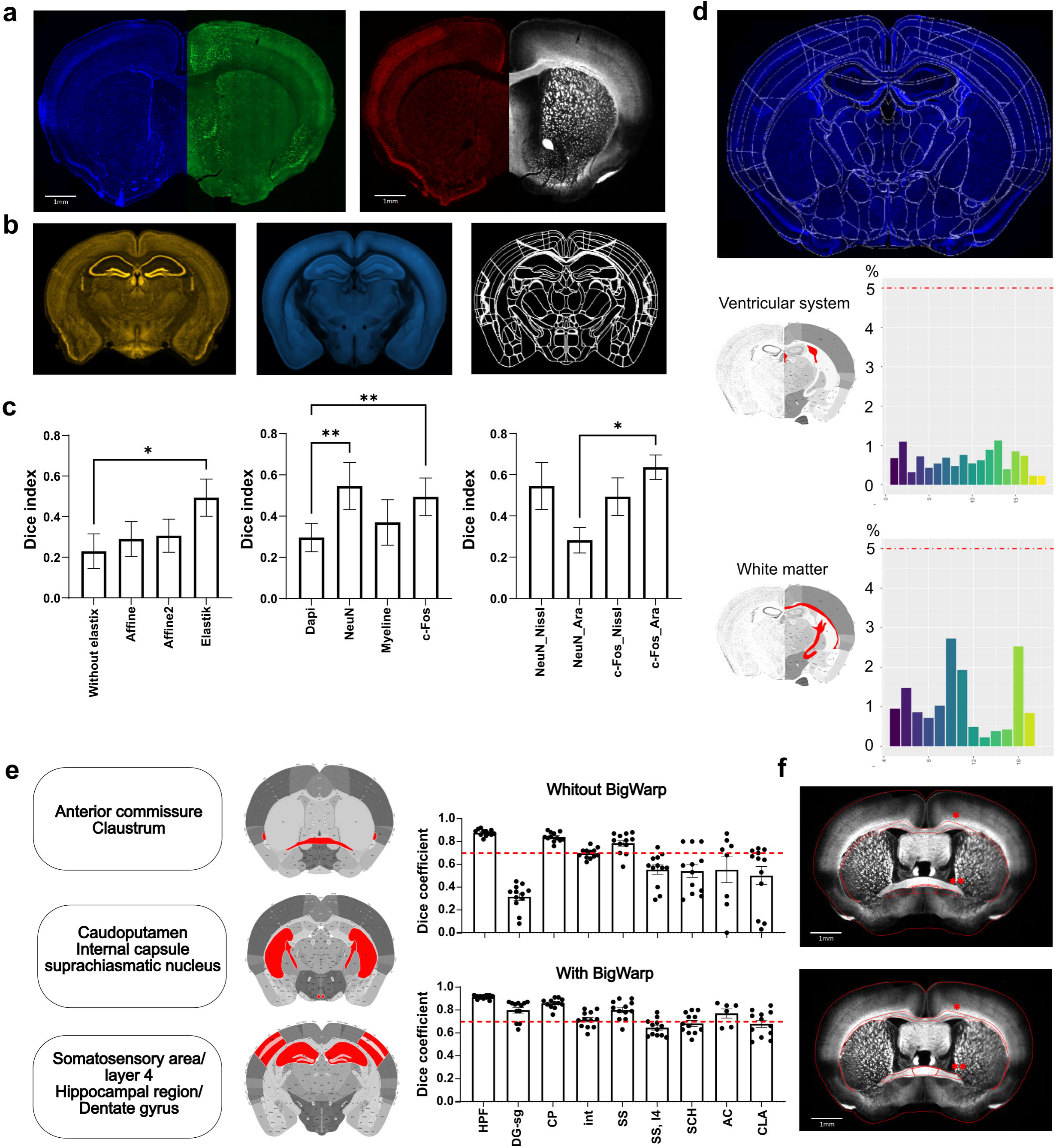
Optimization and validation of atlas registration. **a)** Channel tested for atlas registration: Dapi (blue), c-Fos (green), NeuN (red) and MBP (white). **b)** ABBA atlases tested for atlas registration: Nissl Atlas (brown), Ara Atlas (”3D model”, cyan) and Atlas only composed by brain regions contours. **c)** Comparison of atlas registration accuracy using various channels and/or atlases. Dice correlation for white matter area (n=8) show conditions for which best results are obtained. The first bar graph illustrates the effect of each Elastix algorithm (Affine and Spline) on registration obtained with the c­ Fos channel and the Nissl atlas. Note that Affine and Spline should both be applied for best results, while repeating Affine does not improve accuracy. The second bar graph evaluates the impact of channel selection on registration accuracy using the Nissl Atlas. The final graph shows the impact of choosing different atlases while using the two best channels, i.e. NeuN or c-Fos to perform the registration. d) Regions used (red) for quality control of section registration based on c-Fos detection in regions devoided of neuronal cell bodies, i.e. white matter (WM) and ventricular system (VS). A score higher than 5% indicates misregistration. e) Automatic registration validation by comparison with ROI defined by anatomists. (n=12 per structure). Dice index above 0.7 is considered as good overlap. While registration is optimal for large regions, small and/or tortious regions (i.e. dentate gyrus) necessitate manual refinement using BigWarp. Abbreviations: **HPF:** hippocampal formation; **DG-sg:** Dentate gyrus, granule cell layer; **CP:** caudoputamen; **int:** internal capsule; **SS:** somatosensory areas; **SS,14:** layer 4 of somatosensory areas, **SCH:** suprachiasmatic nucleus; **AC:** anterior commissure; **CLA:** claustrum. **f)** Illustration of registration parameters combination leading to lowest (above, c-Fos_affine_nissl) and highest (below, c-Fos_spline_Ara) Dice index. Red ROI correspond to brain region registration after parameters application (* : Corpus callosum; **: Anterior commissure). Data are expressed as Mean± SEM, * P<0.05, **P<0.01.

Results confirmed the efficiency of sequentially applying linear and nonlinear “elastik algorithms” (p=0.01, One-way Anova, Friedman test) to significantly improve registration accuracy, as evidenced by the Dice coefficient increasing from 0.23 ± 0.08 without anything to 0.49 ± 0.09 when elastik registrations were applied (p<0.05). Our quantification also revealed that repetition of affine registration does not yield further improvement in the Dice coefficient (p>0.99, **Figure 3c**). Further, channel selection also significantly improved registration accuracy (P<0.01, One-way Anova) with a significant Dice coefficient improvement when NeuN and c-Fos channels are used compared to Dapi counterstaining (0.55 ± 0.11 and 0.49 ± 0.09 respectively for NeuN(p=0.0094) and c-Fos(p=0.0076) vs. Dapi 0.30 ± 0.07). Interestingly, MBP counterstaining showed no benefits on registration accuracy. Finally, we aimed at testing reference atlases available on ABBA. Results revealed a more limited impact of the reference atlas selection (p=0.05, One-way Anova), with the 3D model ‘Ara’ resulting in higher registration accuracy using c-Fos channel than NeuN (0.64 ± 0.06 c-Fos_Ara vs 0.28 ± 0.06 NeuN_Ara, p=0.04). Further, dice index obtained with NeuN_Nissl and c-Fos_Nissl show no significant difference with c-Fos_Ara (both p>0.99) providing a certain degree of flexibility for users in atlas selection. Taken together, these findings underscore the importance of selecting the appropriate image channel, reference atlas and registration procedure to consistently achieve the highest Dice coefficient. They further reveal that Dapi counterstaining is inadequate for optimal registration, while myelin counterstaining impedes the process. Based on these results, we adopted elastik transformation (Affine + Spline) in conjunction with the c-Fos channel and 3D model ‘Ara’ atlas for precise section registration in the following analysis.

To systematically validate cell detection and registration accuracy and facilitate identification of aberrant sections, we implemented a quality control step within our workflow. We provide a script (“QuPath_ABBA analysis_QualityControl.R”) that should be used on all sections before proceeding with further analysis. This script can be run on a list of “*csv*” files contained within a folder to automatically generate a figure composed of 6 panels that bring distinct types of information to the experimenter (see **Extended Data Figure 2**). Aberrant numbers should prompt careful consideration to sections, i.e. acquiring new images, excluding them from quantification, or conducting refined analysis. <u>Panels a and b</u> (i.e. *Num.c-Fos/slices* and *density of c-Fos/slices*) facilitate the identification of missing or aberrant sections (e.g. folded or broken sections, out-of-focus sections etc.). <u>Panel c</u> (i.e. *Percentage artifact/slice*), highlights sections presenting aberrant ratios of detected artifacts among the total number of detected cells on all sections. Significant fluctuations in this metric indicate sections containing artifacts such as dust. <u>Panels d and e</u> (i.e. percentage of c-Fos in 2 control regions, i.e. the *ventricular system* and the *white matter* (composed of the *corpus callosum, body, anterior forceps and olfactory limb, the internal capsule,* allowing to cover the entire rostro-caudal extent of the brain) provide important information on registration accuracy, by their absence of c-Fos positive cells (**Figure 3d**). An arbitrary “red line” appears at 5%, corresponding to 5% of detected cells present in those control regions. Sections above this line should receive immediate attention for improving registration by repeating the aforementioned steps or by performing manual adjustments using BigWarp registration refinement step for registration accuracy improvement, as detailed below. Finally, <u>Panel f</u> (i.e. *density of c-Fos in main brain regions*), allows rapidly checking if all main, non-overlapping brain regions are faithfully represented within the dataset. (**Figure 3e**).

We recommend using BigWarp with caution, as excessive use or application to damaged/incomplete sections may introduce local deformations. However, we have devised a workflow that significantly enhances registration efficiency, particularly for small or intricately shaped brain regions (**Figure 3f**). For optimal results we recommend the following steps: 1) realigning section borders when necessary, 2) realigning interhemispheric “midline”, 3) realigning white matter tracts, 4) realigning elongated, sinuous brain regions, such as the dentate gyrus. We validated this workflow by calculating dice coefficients for nine brain regions manually delineated by three anatomists, comparing registrations before and after applying BigWarp. We selected ROIs of different sizes in both dorsal and ventral areas of rostral and caudal sections, i.e. hippocampal formation, dentate gyrus (granular layer), caudoputamen, internal capsule, somatosensory cortex, Layer 4 of the somatosensory cortex, suprachiasmatic nucleus, anterior commissure and Claustrum. Data revealed that Dice coefficient exceeds 0.7 for most medium to large anatomical regions, i.e. somatosensory area (0.80 ± 0.02), caudoputamen (0.86 ± 0.01), hippocampal region (0.91 ± 0.005), with minimal impact from “BigWarp” adjustments. However, registration accuracy for the smallest or more convoluted structures may vary, requiring systematic BigWarp manual adjustment to reach a Dice coefficient equal or exceeding 0.7, e.g., anterior commissure (0.77 ± 0.04, +40% compared to prior-BigWarp), internal capsule (0,71 ± 0,02, +2%), suprachiasmatic nuclei (0.68 ± 0.03, +26%). Together, these results underscore the importance of employing BigWarp for achieving optimal registration of small or intricate brain regions and offer a mean to validate registration accuracy.

### Stage 3- Data extraction, analysis and representation

We refined a selection of brain regions, i.e. “major divisions”, “summary structures’’, “full list” that correspond to distinct hierarchical levels of the 3D Allen Mouse Brain Common Coordinate Framework (CCFv3^14^ ; 9, 206 and 467 regions, respectively, **Figure 4a**). We manually curated these atlases at different resolutions to minimize overlap between brain regions and avoid any gaps within representations. This is important for having a faithful representation of the entire brain at different resolutions, but also to avoid any artifacts when representing results. These atlases facilitate optimal representation of acquired results from BrainRender, a Python library for the visualization of three-dimensional neuro-anatomical data (https://github.com/brainglobe/brainrender). For each region of the selected atlas, the “QuPath_ABBA analysis_Comparison groups.R’’ script will extract c-Fos cell count, the surface of areas and calculate the density or the proportion of c-Fos positive cells per condition. The 3D data representation script will then allow representing c-Fos cell densities within a given condition and/or compare cell densities or proportions between two conditions. Finally, a list of regions exhibiting significant changes between conditions can be generated. A 3D representation of obtained results can be generated with BrainRender, using the “NV_cerveau3D_AllExperiments_v2023-12-12” script (**see Extended Data Figure 3**).

**Fig 4:**
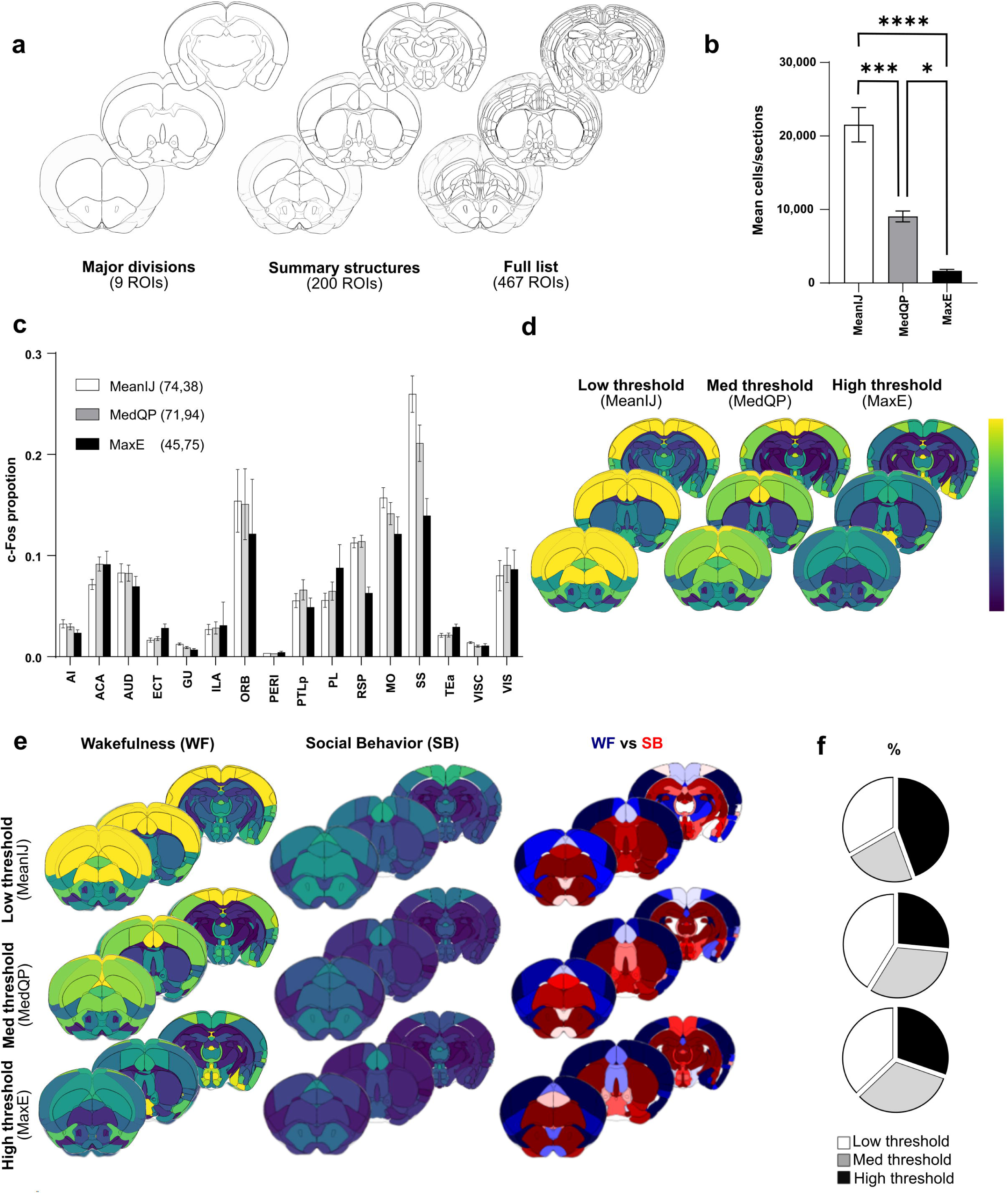
Detection restricted to brighter c-Fos positive cells reduces results relevance. **a)** Illustration of three atlases for whole brain c-Fos analysis at different resolutions. **b)** Automatic thresholding using low (MeanlJ), med (MedQP) and high (MaxE), allows detecting various numbers of cells. **c)** Automatic thresholding methods enable comparison of the proportion of c-Fos positive cells in various brain regions (N=5 mice), here areas composing the isocortex. Note the reduced contrast observed between cortical regions when using higher thresholds, as indicated by values from non-parametric one-way Anova (Krustal-Wallis) statistics showing reduced dispersion of c-Fos proportions. **d)** Corresponding representations of c-Fos cell density at selected rostro-caudal axis coordinates. Note the loss of contrast when using high thresholding methods. **e)** Comparison of c-Fos cell density using “summary structures atlas” in mice (n=5 mice per condition) exposed to two different behavioral paradigms, i.e. Wakefulness (WF) and Social Behavior (SB). A direct comparison highlights region preferentially activated in WF (blue gradient) or SB (red gradient) conditions. **f)** Percentage of regions previously identified as active in social interactions and showing significantly higher activity in SB condition when using Min, Med and High automatic thresholding. Results are shown for “major division” (top), “Summary structures” (middle) and “Full list” (bottom) atlases. Note that high thresholding identifies a lower percentage of relevant regions. Abbreviations: **Al:** Agranular insular area; **ACA:** Anterior Cingulate Area; **AUD:** Audritoy Areas; **ECT:** Ectorhinal Area; **GU:** Gustatory Areas; **ILA** : lnfralimbic Area; **ORB:** Orbital Area; **PERI:** Perihinal Area; **PTLp:** Posterior Partietal Association areas; **PL:** Prelimbic area; **RSP:** Restroplenial area; **MO:** Somatomotor areas; **55:** Somatosensory

Having benchmarked various automatic thresholding methods, which allow unbiased detection of either a large number of cells or a subset (i.e. the brighter cells), we sought to determine the most informative approach for studying brain-wide activity patterns with c-Fos. To this end we selected three levels of c-Fos positive cell detection (**Figure 4b**) and applied them to mice subjected to the “wakefulness” (i.e. WF) paradigm, resulting in different detection levels: MeanIJ, defined as low threshold, (21534 ± 2340 cells/animal); MedQP, defined as med threshold, (9052 ± 747 cells/animal, i.e. 42% of meanIJ), and MaxE, defined as high threshold (1671 ± 186 cells/animal, i.e. 8% of meanIJ). Analysis using the “summary structure” atlas, revealed that higher threshold led to reduced contrast in cell proportion observed in different brain regions, as illustrated for cortical areas (**Figure 4c**). For instance, while most c-Fos cells were detected within the somatosensory cortex of “wakefulness” mice with low and med thresholding methods, this pronounced enrichment was lost when only brighter cells were detected (0.26±0.02 for MeanIJ *vs.* 0.14±0.02 for MaxE, p<0.01). This reduced amplitude of observed differences between cortical areas was supported by a higher K value when using a low threshold (MeanIJ, 74.38; *Krustal Wallis*), gradually decreasing with higher thresholding values (MedQP, 71.94 and MaxE, 45.73). Further, the 3D visualization of c-Fos cell density obtained at the three thresholds clearly highlighted the gradual loss of contrast observed between brain regions **(Figure 4d**). Thus, whereas comparison of low (MeanIJ) and medium thresholds (MedQP) detection led to 94.5% similarities of all ROIs being detected as significantly different, this number decreased to 32% when higher threshold (MaxE) was applied, further supporting a profound reduction of contrast when only brighter c-Fos cells were detected. Noticeably, this effect appeared to be more pronounced within the cortex, as contrast in cell proportion within subcortical structures remained less affected by the methodology used for cell detection (see **Extended Data Figure 4**).

To further analyze the consequences of this reduced contrast for comparisons between different experimental groups, we compared c-Fos positive cell densities in mice subjected to “wakefulness” and “social behavior” paradigms (WF and SB, respectively), using the three thresholding methods outlined above. For a visual inspection of obtained results, we used a color code highlighting differences between conditions, i.e. a blue gradient associated with regions activated in WF and a red gradient associated with those activated in SB (**Figure 4e**). Regions showing significantly higher levels of activity in “social behavior” were listed at all hierarchical atlas levels (“major divisions”, “summary structures’’, “full list”), and compared to those previously identified in a classical study (^15^, **Figure 4f**). While both MeanIJ and MedQP thresholding consistently yielded comparable outcomes in identifying regions previously identified as activated during social behavior, results obtained with MaxE showed more mitigated results.

In summary, our careful benchmarking underscores the importance of integrating all pre-established optimal parameters for a thorough whole-brain analysis. Specifically, we recommend using MeanIJ or MedQP thresholding methods alongside a “large sample size” classifier refinement to ensure accurate detection of condition-specific neuronal activity patterns. Furthermore, for ABBA atlas registration, we suggest implementing recommend employing Elastik registration using c-Fos/NeuN markers in combination with Ara or Nissl atlases, followed by refinement through BigWarp registration for optimal results.

### Identification of distinct activity patterns in different behaviors and at different scales

We began by validating our workflow by comparing c-Fos expression patterns in groups of mice (n=5) subjected to distinct behavioral paradigms, i.e. Wakefulness (WF), Social Behaviour (SB) and Paradoxical Sleep (PS). Analysis of c-Fos positive cell densities within each group revealed distinct activity patterns among the three conditions (**Figure 5a**). For example, activity in somatosensory and motor cortical areas were prevalent in WF, while activity in frontal cortical areas and in periventricular regions were more pronounced in SB and PS, respectively (**Figure 5a**). These differences were however accompanied by pronounced differences in activity levels between groups, with the number of c-Fos positive cells being significantly lower in PS compared to WF or SB (Suppl **Figure 5a-b**). This discrepancy prompted us to investigate whether such substantial differences in activity might hinder the detection of condition-specific patterns of c-Fos expression. To address this, we first compared results obtained by quantifying raw cell numbers and cell density within the cortex (**Extended Data Figure 5c, d & 6a, b**). As anticipated, comparison of SB or PS with WF revealed pronounced effects of activity levels on obtained outcomes. For instance, all cortical regions appeared to be significantly lower in PS/SB than WF, reflecting the global decrease of activity while impeding the identification of condition-specific cortical activity patterns.

**Fig 5:**
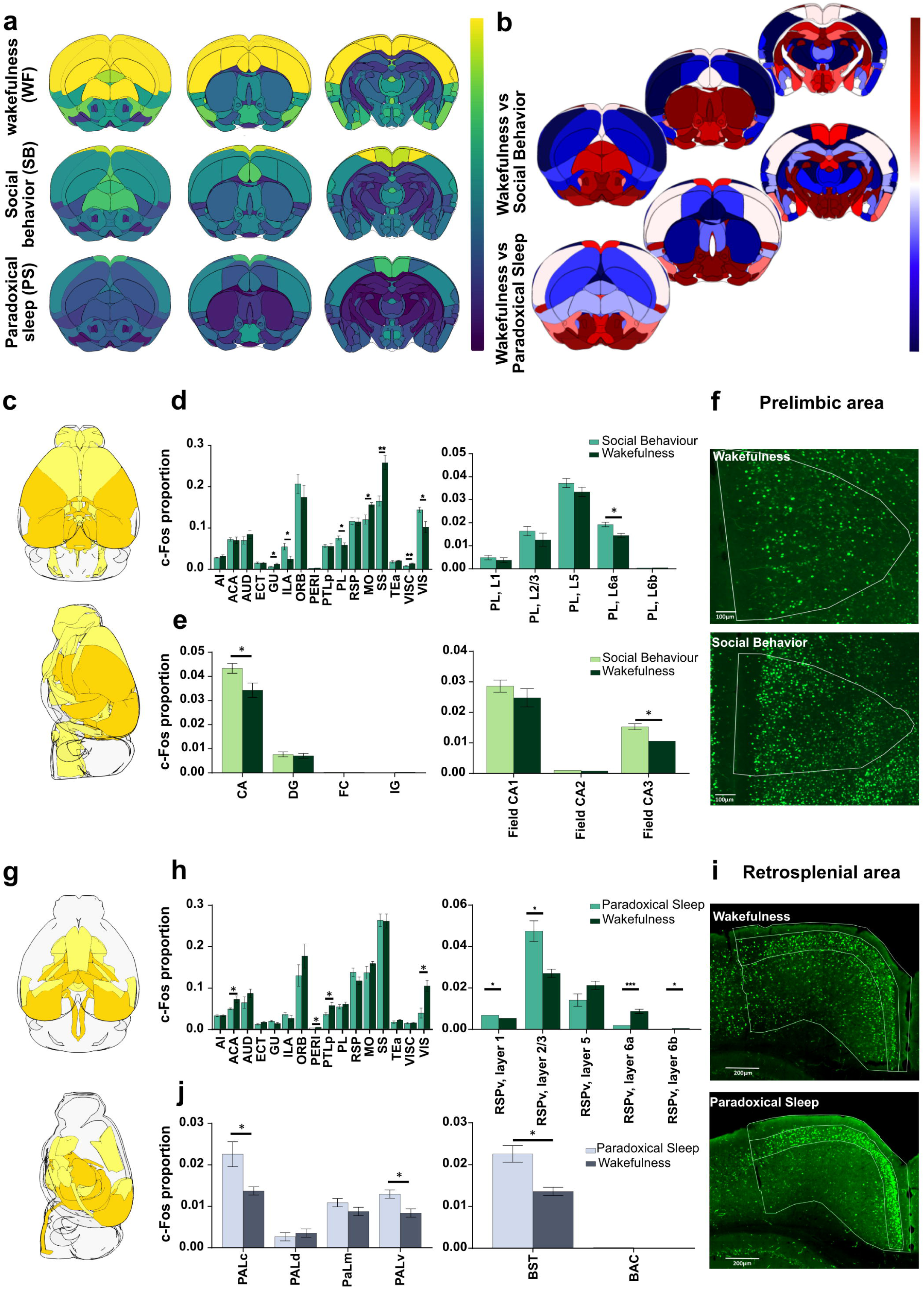
Spatial analysis of activity patterns in different behaviors and at different scales. c-Fos expression patterns related to different behaviors can be visualized and explored using several qualitative and quantitative representations. **a)** 3D projections of c-Fos density in various brain regions (”Summary structure” atlas). **b)** Direct comparison highlighting regions preferentially activated in WF (control condition, blue gradient), when compared to SB or PS (red gradient, upper and lower sections respectively). **c-i)** Quantitative data analysis can be visualized by two methods. First, by identification of brain regions showing significant differences in c-Fos proportions on a 3D model, i.e. here for lsocortex areas **(c, g,** light yellow = P<0.05; dark yellow= P<0.01). Second, by using bar graphs for defined brain regions at different resolutions (”summary structures”, left, “full list”, right)). Analysis of the lsocortex **(d, h)** shows different patterns of c-Fos expression following SB and PS. Focus on specific isocortical areas, i.e. prelimbic area **(d)** or retrosplenial area **(h)** highlight more discrete differences. **e, i)** Corresponding immunostainings illustrating global **(f)** or layer specific **(i)** c-Fos induction. **e, j)** Analysis of the hippocampal formation and pallidum at similar increasing resolution (i.e ammon’s horn and pallidum medial/caudal region, respectively) highlight significant differences in c-Fos expression pattern in SB **(e)** and PS **(j)** conditions. Abbreviations: **Al:** Agranular insular area; **ACA:** Anterior Cingulate Area; **AUD:** Auditory Areas; **ECT:** Ectorhinal Area; **GU:** Gustatory Areas; **ILA** : lnfralimbic Area; **ORB:** Orbital Area; **PERI:** Perirhinal Area; **PTLp:** Posterior Parietal Association areas; **PL:** Prelimbic area; **RSP:** Restroplenial area; **MO:** Somatomotor areas; **SS:** Somatosensory areas; **TEa:** Temporal Association areas; **VISC:** Visceral area; **VIS:** Visual areas; **PAL** : Pallidum **(d** : dorsal, **c:** caudal, **m:** medial, **v:** ventral); **BST:** Bed nucleus of stria terminalis; **BAC** Bed nucleus of the anterior commissure. Data are expressed as Mean ± SEM, *P<0.05; ** P<0.01

To facilitate extraction of these nuanced changes in c-Fos expression patterns between conditions, we devised a method to calculate a c-Fos positive cells proportion per brain region, by normalizing cell counts with the total number of cells detected per section. For qualitative visual representation of obtained results, we used a color gradient highlighting differences between conditions, i.e. a blue gradient indicating regions activated in WF and a red gradient indicating those activated during SB or PS (**Figure 5b, Extended Data Figure 5e**). We complemented these visual representations by more quantitative analyses that can be automatically performed at different scales. To this end, we incorporated calculation of p-values between conditions in our script, allowing their representation on 3D brain models (yellow = p<0.05, orange = p<0.01, **Figure 5c-g, Extended Data Figure 6c**), alongside to bar graphs (**Figure 5d-e, h-j**). Notably, this analysis can be performed at different resolutions, by simply selecting appropriate atlases.

To illustrate the versatility of this approach, we compared the c-Fos labeling patterns within cortical areas between WF and SB (**Figure 5d-f**) or WF and PS mice (**Figure 5h-j**), which we complemented by performing a more detailed analysis of cortical layers in select areas. Using this multiscale approach, our results revealed significantly increased c-Fos expression is SB compared to WF mice in the infralimbic (0.025 ± 0.007 for WF and 0.055 ± 0.008 for SB ; p=0.03) and prelimbic cortices (0.060 ± 0.005 for WF and 0.075 ± 0.007 for SB ; p< 0.05), while other regions such as the retrosplenial area showed no significant changes (**Figure 5d**). Focusing on the prelimbic area, our results refined these initial observations by highlighting that the observed increase in c-Fos expression was primarily localized within layer 6a (**Figure 5d**). Similarly, our results showed an increased c-Fos proportion within the Ammon’s horn of the hippocampal formation in SB mice (0.034 ± 0.003 for WF and 0.043 ± 0.002 for SB ; p=0.04), which analysis at higher resolution demonstrated to be specific to the CA3 fields (CA3: 0.010 ± 0.000 for WF and 0.015 ± 0.001 for SB; p=0.02, **Figure 5e**). Finally, while we found a higher c-Fos proportion during PS in the caudal and ventral pallidum, analysis at higher resolution refined those observations by showing involvement of the bed nucleus of stria terminalis (0.022 ± 0.002 for PS and 0.014 ± 0.001 for WF, p=0.02, **Figure 5j**). Such multiscale analysis also enabled the identification of layer-specific changes in c-Fos expression patterns, which were not evident when considering the cortical area as a whole. For instance, while no significant difference in c-Fos proportion was noted within the retrosplenial area when comparing WF and PS mice **(Figure 5h)**, a more detailed analysis highlighted a redistribution of c-Fos expression towards layers 2/3 during PS (0.027 ± 0.002 for WF and 0.047 ± 0.005 for PS; p=0.02 for ventral part, see also immunostainings in **Figure 5j**).

Altogether, these results underscore the pertinence of comparing c-Fos positive cells raw numbers or density to reveal changes in behavior-related activity levels, while the calculation of c-Fos cell proportions enables the identification of nuanced changes in activity patterns. Further, our results emphasize the importance of conducting analyses at various anatomical resolutions, a principle which was therefore fully implemented within provided scripts.

### Application of the automated workflow to compare activity induced by distinct behaviors in the TRAP2 mice

While previous analyses were performed in distinct animal groups, transgenic TRAP2 mice allow comparing behaviorally-induced activity patterns within the same anima⍰^6^;^17^. These mice express an inducible CRE-recombinase driven by c-Fos promoter leading to the permanent expression of tdTomato upon administration of 4-hydroxytamoxifen (OH-Tam) (**Figure 6a**). Consequently, this transgenic mouse model enables the comparison of activity patterns elicited by different behaviors within individual animals. Here, we initially subjected transgenic TRAP2 mice to a wake procedure 120 minutes prior to OH-Tam injection. Subsequently, the same animals were exposed a week later to a PS deprivation and rebound and were sacrificed 120 minutes later for c-Fos immunodetection (**Figure 6b**).

**Fig 6:**
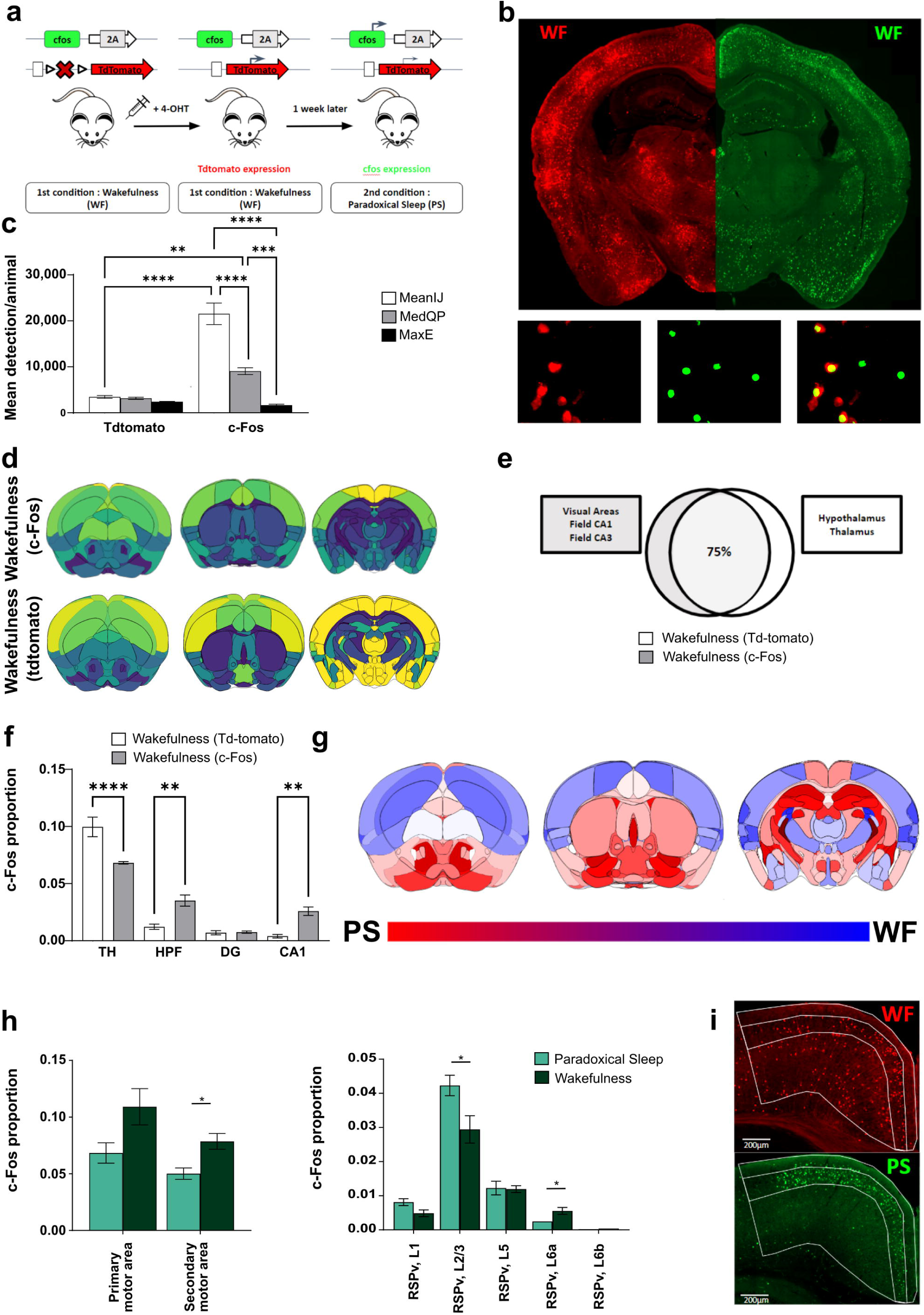
Temporal Analysis of Activity Patterns Using TRAP2 Mice. **a)** Workflow for experiments with TRAP2 mice. 4-OHT (80 mg/Kg, i.p.) was injected 120 minutes following a wakefulness (WF) task to induced c-Fos-driven tdTomato expression (red signal). One week later, mice were exposed to paradoxical sleep (PS) and perfused 120 minutes later for c-Fos immunodetection (green signal). **b)** Illustrative hemisections showing tdTomato expression and c-Fos immunodetection and corresponding automatic detections. **c)** Quantification of mean number of detection per animal using low (MeanlJ), med (MedQP) and high (MaxE) thresholding methods for tdTomato-positive cells and c-Fos-positive cells. **d)** 3D projections of c-Fos and tdTomato density in wakefulness. **e)** Diagram representing the percentage of similarity observed in c-Fos and tdTomato pattern of expression in Wakefulness. While most (i.e. 75%) activated regions are common between both models, some are significantly more activated following c-Fos immunodetection (gray box) or tdTomato induction (white box). **f)** Proportion of tdTomato and c-Fos immunodetection in various subcortical structures. **g)** Quantitative 3D projections of results obtained with tdTomato mice, when comparing wakefulness (WF) vs. Paradoxical Sleep (PS)-induced activity (blue and red gradients represent structures activated during W and PS, respectively). **h)** Comparison of Somatomotor and Retrosplenial Area (RSP) activity on PS/WF conditions. **i)** c­ Fos/tdTomato immunostaining comparing RSP sub-structures activity on WF(tdTomato, red) and PS(c-Fos, green). (N=4 mice per condition). Abbreviations: **RSPv:** retrosplenial area ventral part; **HPF:** Hippocampal formation; **DG:** Dentate gyrus; **TH:** Thalamus. Data are expressed as Mean ± SEM, *P<0.05; **P<0.01; ***P<0.001.

We first compared the number of detected tdTomato-positive cells using our three automatic thresholding methods. In contrast to results obtained for c-Fos immunostaining (i.e. MeanIJ, MedQP, MaxE; **Figure 6c)**, only limited differences were observed in the number of detected tdTomato-positive cells, illustrating the rather homogeneous reporter gene expression. As a consequence, the ratio of c-Fos positive cells detected by immunostaining vs tdTom expression varies significantly when using various automatic thresholding methods. For instance, tdTomato positive cells represented ∼1/6th of the number of detected c-Fos immunoreactive cells when using MeanIJ thresholding methods, ∼1/3rd for MedianQP and an equal number for MaxE thresholding method (**Figure 6c**).

We proceeded to compare the distribution of positive cells induced by wakefulness using c-Fos immunostaining or tdTomato detection. Visualization of cell density on a 3D model using the “Summary Structures Atlas’’ revealed a largely similar pattern of activity between c-Fos and tdTomato detection. However, closer inspection reveals a higher density of tdTomato+ cells in the diencephalon when compared to c-Fos immunostaining (**Figure 6d**), suggestive of a higher expression of cFos and therefore a higher expression of CreERT2 and resulting tdTomato induction in subcortical areas. Direct comparison of cell distribution between the two groups of mice (**Figure 6e**) confirmed that 75% of the regions exhibited similar activity levels. For the remaining 25%, most regions showing higher activity following c-Fos immunostaining were related to the isocortex, i.e. visual areas and Field CA1/CA3 (i.e. 0.070 ± 0.013 for tdTomato vs 0.126 ± 0.017 for c-Fos immunodetection, P< 0.001). Conversely, regions with higher activity in tdTomato mice were primarily related to the thalamus and hypothalamus (i.e. 0.09 ± 0.08 for tdTomato vs 0.06 ± 0.004 for WF c-Fos on Hypothalamus, P< 0.01, unpaired T-test). These observations were further supported by comparing cell density within the thalamus, hippocampal formation, dentate gyrus and CA1 subregions, in both groups of mice (**Figure 6f**). Comparison of positive cell proportions revealed higher activity in the thalamus using tdTomato detection (i.e. 0.10 ± 0.008 for tdTomato *vs* 0.07 ± 0.001 for c-Fos, p<0.01), while higher activity was observed within the hippocampal formation and CA1 when using c-Fos immunodetection (i.e.0.004 ± 0.001 for tdTomato *vs* 0.0259 ± 0.004 for c-Fos in CA1, p<0.01). Thus, although c-Fos immunodetection and tdTomato induction show largely similar patterns of activity during wakefulness, the former appears to be more sensitive for cortical regions, while the other shows greater sensitivity within subcortical regions.

We then sought to investigate whether these differences could affect results obtained when comparing activity patterns induced by distinct behaviors, here WF and PS. We compared results obtained when using c-Fos immunostaining in separate groups of mice (see **Figure 5**), to those obtained in TRAP2 mice sequentially exposed to the same two behavioral paradigms (**Figure 6g**). Qualitative 3D representation revealed pronounced similarities in the pattern of activation induced by both behaviors when using both approaches (**Figure 6g**). As tdTomato may underestimate activity within the cortex, we next focused our analysis onto this region. Our analysis in TRAP2 mice replicated most of the results obtained in separate groups of mice (WF and PS groups). Notably, a similar increase in activity was observed in the secondary motor area during WF, albeit significantly higher with tdTomato detection when compared to c-Fos staining (0.078 ± 0.007 WF *vs* 0.050 ± 0.005 PS, p= 0.02; **Figure 6h**). Additionally, for regions like the retrosplenial area, a higher-resolution analysis confirmed elevated activity in layer 2/3 during PS, when compared to WF (0.042 ± 0.003 for PS vs 0.029 ± 0.004 for WF, P =0.04), as observed using c-Fos staining.

Taken together, our results indicate that the workflow presented here can be used with TRAP2 mice to enable a direct comparison of activity patterns induced by distinct behaviors within the same group of mice. Importantly, while regional differences may marginally influence obtained results, the overall utility and applicability of the approach remain robust.,

## DISCUSSION

In this study, we established and validated a workflow for semi-automatic whole-brain quantification of single or multiple markers distribution. Our approach offers a complementary alternative to existing methods that often require specialized equipment (e.g., STP tomography^18^), or sophisticated analytical platforms (e.g., transparisation^19^).

While several other workflows have been proposed recently, they often rely on commercial software (e.g. Neuroinfo^20^) or on alternative software, which in our hand required higher computer resources (e.g. Quint^7^) than selected ones. Here we took care to select software that are open source and continuously updated, including QuPath for cell segmentation and ABBA for sections registration onto a reference atlas. Notably, although we primarily demonstrate the efficiency of our workflow for quantifying nuclear markers like c-Fos, our quantifications in TRAP2 mice indicate its accuracy for cytoplasmic staining such as tdTomato-induced expression in the Ai14 reporter mouse. We believe that this adaptable workflow can be easily customized for various staining applications, even those involving complex cell types with intricate morphologies. For instance, QuPath has been recently successfully used for quantifying microglial cells, a cell type which exhibits diverse complex morphologies within the central nervous system^21^. Thus, our proposed workflow offers versatility across a range of applications with minimal modifications to the provided scripts. Further, the ongoing development of a multiplex analysis workflow in Qupath holds promise for enabling complex multi-channel quantifications^22,23^ using the methodology outlined here.

Although the section registration workflow using ABBA is comprehensively described on its companion website (https://abba-documentation.readthedocs.io/), we provide users with recommendations as well as quantifications for comparison of outcomes obtained using various combinations of channels and elastic transformations. Alternatives to the proposed manual registration workflow have recently emerged, such as DeepSlice^24^, accessible through an online interface (www.DeepSlice.org) with drag-and-drop functionality. Our testing of this fully automatic approach yielded variable results, with some sections showing registration precision comparable to that obtained with our workflow, while others from the same animal were misregistered (i.e. placed at suboptimal rostro-caudal locations). It is likely that training DeepSlice with sections deposited by users will rapidly enhance the robustness of its performance in the near future. Further, future integration of DeepSlice in the described workflow will be facilitated by its availability as an open-source Python package (github.com/PolarBean/DeepSlice).

The processing of large numbers of sections relies heavily on automation. To address this, we have developed scripts that enable continuous quality control and refinement at three key stages of the workflow. First, we developed an interaction between QuPath and ImageJ for the automatic thresholding of cell detection. This feature allows users to use various thresholding approaches, within QuPath or ImageJ (e.g. Otsu method^25^ for unbiased threshold detection). Approaches such as Cellpose and Stardisk, available within Qupath, may offer alternatives to automatic cell detection presented here, but were not systematically compared. Although efficient, these algorithms do not allow to automatically define different detection thresholds but require a learning file based manual detections produced by users, possibly inducing experimenter bias. Our approach, based on automatic thresholding minimizes detection variability between sections, resulting in detections of cells with defined c-Fos staining intensities that appear to be of biological significance (see below). Second, we included in this workflow principle for classifier refinement using machine learning to allow efficient removal of blood vessels and/or staining artifacts. This is key in maximizing the number of exploitable sections/animals despite variability in perfusion efficiency, aligning with the 3Rs principles. Third, we have meticulously benchmarked the section registration workflow and developed scripts for quality control. These tools allow users to identify suboptimal sections and refine registration either automatically or manually using BigWarp. Finally, we provide a well annotated script for comparison and visualization of results obtained in different experimental groups. This comprehensive approach ensures the robustness and reproducibility of the analysis across large datasets, facilitating meaningful interpretations of experimental outcomes.

A recurrent challenge in c-Fos quantifications revolves around selecting an appropriate threshold to ensure that only cells relevant to the behavioral task are counted, while discarding irrelevant background signal. The c-Fos protein starts to be produced approximately 15 minutes after stimulation, and is therefore named an immediate early gene. It is expressed in both excitatory and inhibitory neurons^26,27^, and serves as an indirect marker of neuronal activity, as its expression is transient but contingent upon neurons generating action potentials^28^. Because of these dynamic properties, c-Fos staining results in a spectrum of intensities across various neurons. As c-Fos expression is transient, it is conceivable that low-expressing cells represent cells activated before or after animals were exposed to a given stimulus. Alternatively, it is also possible that the level of c-Fos expression is different in distinct subtypes of neurons, with some publications reporting low signal in glial cells^29^. Previous studies in a PTZ-induced seizure rat model revealed transient c-Fos mRNA expression, peaking 30 minutes following seizure induction and declining after 1 hour to return to baseline 6 hours post-seizure^30^. Determining the peak of c-Fos expression is more challenging when testing animals in complex, protacted behavioral tasks, such as those used in our study. Conventionally, a survival time of 1h30 following task completion is used, and selection of the most significant signal is achieved by adjusting the detection threshold of c-Fos positive cells. Here, we show that automatic thresholding, alongside minimizing intersection detection variability, also facilitates detecting cells exhibiting various c-Fos staining intensities. Surprisingly, our results suggest that most intensely c-Fos positive neurons may not necessarily be the most relevant to the observed behavior-induced neuronal activity. Indeed, detection of a larger number of cells, including those with dim labeling, enhances the discrimination between patterns of behaviorally-induced activity. This is in line with mathematical approaches comparing differences between control and experimental groups, which have defined that ∼60% of the most positive neurons have to be quantified to observe maximal differences between groups^30^.

It is noteworthy that presenting results as raw cell numbers, cell densities or cell proportions, conveys distinct types of information. While quantifying cell numbers or cell density between conditions provides essential insights into activity levels, expressing results in proportion allows for a deeper exploration of task-related activity patterns. This distinction became particularly evident when comparing behaviors eliciting varying levels of activity, such as WF and PS. For these two different behaviors, the number of c-Fos positive cells in WF mice vastly outnumbered those of PS mice, therefore hindering crucial insights into the respective distribution of these cells. To address this limitation, we developed a normalization method based on “proportion” calculation, enabling a more direct comparison of cell distribution during specific behaviors. Given the complementarity of these analyses, we have implemented all three quantification methods into the provided scripts.

Using thresholding approaches allowing detecting > 50% of c-Fos positive neurons, we identified activity patterns in both WF/SB and WF/PS paradigms that are in agreement with existing literature. Thus, structures predominantly activated during social behavior included the prefrontal cortex (specifically, the prelimbic and infralimbic areas) as previously showed^31–33^, as well as the CA3 hippocampal regions, which are known for their role in memory storage and social recognition memory ^34,35^. Moreover, our comparison of structures activated during wakefulness and paradoxical sleep highlighted predominant activity in the anterior cingulate during WF, while the dentate gyrus exhibited increased activation during PS, consistent with previous reports^36^. Of note, our approach enabled the detection of more discrete subregions, including cortical layers, confirming the activation of retrosplenial area layer 2/3 neurons during PS^37,38^, thereby illustrating the workflow’s capability to detect changes in activity at various scales, from brain regions to subregions/layers. The capacity to perform this analysis at the whole brain level further unveiled activation of structures not traditionally associated with the studied behaviors. For instance, we observed a significant increase in c-Fos expression in the Bed nucleus of stria terminalis (BNST) in PS mice. The BNST is part of the extended amygdala, a structure implicated in stress response, fear and anxiety. While global activation of GABA neurons in the BNST during slow wave sleep^39^ induces wakefulness, it does not rule out that subpopulations of cells in this nucleus might play a specific role in PS. For such newly identified brain regions, further investigations such as connectivity analysis may be used to better understand their role in specific behavioral tasks ^40–42^.

Our workflow can also be used in animals sequentially exposed to different behavioral tasks. To explore such capability, we used TRAP2 mice^17^, which allows the induction and permanent expression of a reporter gene in c-Fos-expressing neurons upon tamoxifen injection. For this experiment, we separated behavioral tasks by a week. Indeed, after a first stimulus c-Fos expression has been reported to show refractory period to subsequent stimulation of similar nature^30^, thereby possibly masking reactivation of neurons. In rodents, this refractory period is short, i.e. 3 to 6 hours. This suggest that the temporal exposure of the same mice to subsequent behavioral tasks may be reduced. Our results demonstrate that largely similar results were obtained when using TRAP2 mice compared to staining performed in separate groups of mice. However, small discrepancies were observed, which may be attributed to two factors: firstly, the longer half-life of the cre-recombinase protein when compared to the transient c-Fos expression, and secondly variation in the level of cFos expression observed across brain regions.

Collectively, our workflow allows generating high-resolution maps of behaviorally-evoked whole-brain activation in mice. Its automation, combined with the integration of quality control steps throughout the workflow, allows for optimization for different experimental setups and c-Fos detection methods. By analyzing large numbers of sections, our approach enables the comparison of activity patterns across brain regions, facilitating unbiased whole-brain analysis and the identification of novel structures activated during specific tasks. Moreover, this versatile workflow can be applied to rodent models of neurological diseases and disorders to uncover circuit deficits associated with alterations in social, cognitive, and other higher-order brain functions.

## MATERIAL AND METHODS

### Animal

five OF1 mice (Charles River, France) were used for the project. Normoxic experiments were performed according to European requirements 2010/63/UE and have been approved by the Animal Care and Use Committee CELYNE (APAFIS #18355, APAFIS#21351).

Ten TRAP2 mice were also used. Male double heterozygous Fos^2A-iCreER^;R26^Ai14^ (TRAP2^16^) mice were used in paradoxical sleep and wake experiments, generated by crossing Fos^2A-iCreER/+^ (TRAP2) mice to R26^AI14^/+ (AI14) mice. TRAP2 mice were kindly gifted by Dr Liqun Luo from Stanford University.

For all experiments, age 8-12 weeks old mice were used. Mice were housed individually and placed under a constant light/dark cycle (light on from 8:00 am to 8:00 pm), with *ad libitum* access to food and water.

### Behavioral tests

Mice (N=5 mice per group) were exposed to defined behavioral paradigms, then sacrificed 90- 120 following test completion.

#### Social Behavior (SB)

the social behavior test consisted in a simplified version of the three-chambered social test^43^. In this test, each mouse followed a habituation period for 20 minutes in a plexiglass chamber with a round wired compartment placed in its center. After the habituation period, a new mouse was placed in the round wired compartment for 10 minutes.

#### Wakefulness procedure (WF)

The wakefulness system consists of a white area (40*40 cm) delimited by 4 walls. During the protocol, mice were placed in the box together with wood tips and small objects to maintain a continuous awakening for 4 hours. Animals were permanently monitored by the camera from a different room to check whether the animals were awake. When the mouse was inactive/drowsy, it was gently touched by the experimentator and objects were moved.

#### Paradoxical Sleep (PS)

In Order to evaluate the pattern of neuronal activation during PS, 2h of PS hypersomnia was induced by exposing the mice to 48h of PS deprivation. To this end, electrodes were implanted for automatic detection of PS episodes, as previously described^44,45^. Briefly, mice were anesthetized using a cocktail of ketamine and xylazine (100/10 mg/kg; Imalgene 1000, Merial and Rompun 2% Bayer) and 4 EEG electrodes and 2 EMG were implanted^38^. Following 7 to 10 days recovery, mice’s vigilance state (Wakefulness, slow-wave sleep, Paradoxical sleep) were identified by visual inspection of polysomnographic signals according to the criteria described in detail previously^44,45^. For PS deprivation, mice were individually placed in a transparent barrel with a movable floor connected to a small piston controlled by an electromagnet. The automatic detection of PS episodes was achieved by means of a learning file based on the parameters of the sleep-wake cycle states previously defined during baseline recordings. PS deprivations started at 10:00 am. During the recordings, PS was detected computationally and in real time by an algorithm, a digital transistor-transistor logic pulse was sent to a current generator controlling the instantaneous onset of the electro-magnet. It caused movements (up/down) of the floor, resulting in a waking up of the animal after only 5 s from the detection of a PS^45^. After 48 h of PS deprivation, the system automatically stopped at 10:00 a.m. Then animals were allowed to recover and to display PS hypersomnia in the same barrel.

### Tissue preparation and acquisition

All mice were sacrificed by an intraperitoneal overdose of pentobarbital followed by transcardial perfusion with Ringer Lactate solution followed by 4% paraformaldehyde (PFA) dissolved in 0.1□M phosphate buffer (PB; pH 7.4). Brains were rapidly removed and postfixed for 48 hours in 4% PFA at 4□°C. For cryostat sectioning (see below), the brains were immersed in a 30% Sucrose 0.1M PB solution for 72 hours before being frozen in 2-methylbutane placed on dry ice at around −33°C.

#### Brain sectioning

Brains were cut into sections of 50µm (SB) or 30µm (WF and PS) of thickness using a vibratome (VT1000 S; Leica; Wetzlar; Germany) or a cryostat (Leica CM1950), respectively. Series of 12 (SB) or 8 (WF and PS) sections were collected in 12 wells plate and preserved at −20°C in cryoprotective solution containing 20% glycerol and 30% ethylene glycol in 0.05 M PB (pH 7.4).

For immunochemistry, 2 wells (series of 1:6 for SB and series of 1:8 for WF and PS) were used, to ensure homogeneous sampling of the ROI and sufficient number of sections per structure according to stereological standards.

#### Immunohistochemistry

Brain sections were first washed in 0.1 M PBS and 0.4% Triton X-100 (Sigma-Aldrich) to remove the cryoprotectant. Sections were then incubated in 0.3% H2O2 for 1 hour to quench endogenous peroxidase activity, then washed 3*10 minutes in 0.1 M PBS and 0.4% Triton X-100. All sections were then incubated with anti-c-Fos rat antibodies (c-Fos antibody - 226 017; Synaptic System, 1:50000) for 48 h at 4°C in 0.1 M PBS 0.4% Triton X-100 containing, then washed 3*10 minutes in 0.1 M PBS and 0.4% Triton X-100. Sections were then incubated for 2 hours in 0.1 M PBS and 0.4% Triton X-100 containing biotinylated rabbit Anti-Rat IgG antibody diluted to 1:1000 (Vector Laboratories) and washed 3*10 minutes with 0.1 M PBS and 0.4% Triton X-100. Following an incubation of 2 hours with streptavidin (SA)-HRP (Alexa Fluor™ Tyramide SuperBoost™ Kit, streptavidin; Life Technologies, 1:1000) in 0.1 M PBS and 0.4% Triton X-100, sections were washed 3*10 minutes in 0.1 M PBS and 0.4% Triton X-100, then incubated for 10 min in Alexa Fluor 488-conjugated tyramide (Molecular Probes, Eugene, OR, USA) by diluting the stock solution 1:500 in 0.0015% H2O2/amplification buffer. This reaction was terminated after 10 min by rinsing the tissue in 0.1 M PBS. Finally, sections were incubated for 5 minutes in 4′,6-diamidino-2-phenylindole diluted to 1:5000 (1µg/ml) in 0.1M PB solution, then mounted onto gelatin-coated slides before cover slipping.

#### Image acquisition

Sections were imaged using an Axioscan slide scanning microscope (Zeiss, Germany). Images were collected with a x20 objective (N.A. Plan-Apochromat 20x/0.8 M27 Air/0.8.) and a 0.45 Orca Flash camera. Dapi, c-Fos (alexa-488), tdTomato, and NeuN or MBP (alexa-633) were acquired using appropriate filter cubes, according to manufacturer recommendations. Exposure time was defined by signal distribution across gray values (>20% of measure gray values on 16 bits images). For all sections, five mosaics were acquired (5um steps) then projected on a single plane, using the depth of focus “wavelet” function of the Zen software (version 3.1) Images were not modified and were directly imported in QuPath for quantifications.

### c-Fos immunoreactivity analysis

#### Automatic thresholding

We developed an approach allowing automatic adaptive thresholding within QuPath or by using ImageJ thresholding methods. Within QuPath (*version 0.4.3*), individual sections were delineated in order to extract the Median Intensity value. For each section, this value was then transferred to detection parameters in order to set up the level of threshold used for c-Fos cell detection. This procedure can be used by downloading the script *“*Automatic threshold_application_(QuPath)*”*. For using ImageJ thresholding methods (“Otsu” and “Huang” methods for exemple^46^, we developed an additional script *“*Automatic_threshold_application_(ImageJ)*”*. *For further description on these scripts please see below on ‘’Provide scripts’’ section)*.

#### Atlas registration

ABBA software allows section registration on a reference Atlas as well as an optimal communication with the QuPath software. For optimal section registration, we have established a workflow complementing indications provided by the software editor (see https://abba-documentation.readthedocs.io/). Entire description of the workflow is available in the results and “detailed procedure” parts of this manuscript. For final analyses, elastic transformations (affine and spline) were performed using the c-Fos fluorescent channel coupled to the Ara reference Atlas (3D model-like). 10 “control points along X ‘’ were used during the spline step. BigWarp manual registration refinements were achieved with association of c-Fos and Dapi channel, coupled to “label borders” reference atlas.

#### Data Analysis

All analyses were performed on R version 4.3., using RStudio 2023.09.1 Build 494.

### Provided scripts

A description of all scripts can be found on the github page: https://github.com/sebastien-cabrera/Scripts-for-quantification-and-analysis-CABRERA-et-al.

### Statistical analysis

Data obtained were analyzed in R, then validated in GraphPad Prism. Normality distribution was assessed by using the Shapiro-Wilk test. When the assumption of sphericity had been violated, data were analyzed using analysis of variance (ANOVA) corrected by the Greenhouse–Geisser method.

For variability analysis of automatic thresholding methods, we used repeated measures one-way ANOVA followed by a Dunnett’s multiple comparison test. For variability percentage analysis on heterogeneous and homogeneous on consecutive sections, we used Kruskall-Wallis test followed by Dunn’s multiple comparisons test. For the comparison of two “training” methods to distinct artifacts and positive cells by classifier with “small sample size” and “large sample size”, we used two-way anova. For the analysis of the atlas registration accuracy by Dice coefficient, to compare the efficiency of elastix algorithm, channel selected and atlas comparison, we used Friedman test followed by Dunn’s multiple comparisons test. For comparison of detection levels in wakefulness mice using different thresholding methods, we used one way anova test followed by Tukey’s multiple comparison test. For amplitude analysis of low, med and high threshold we used the Kustral Wallis test. For analysis and identification of brain regions activity patterns in different behaviors with 2 cohorts of animals and genetical TRAP2 model, we used parametric unpaired t-test. For comparison of tdTomato and c-Fos positive cells detection following three thresholding methods, we used one-way anova followed by Tukey’s multiple comparison test. For proportion analysis of tdTomato and c-Fos positive cells in subcortical structures we used one-way anova followed by Sidak’s multiple comparison test. Differences were considered significant when P < 0.05.

## Detailed procedure

1. **Create a project in QuPath** □ **Timing**: 1 min

a. Open QuPath software and select “Create Project” by defining folder location
b. “Add images’ to process within the project and choose the appropriate “image type” (e.g. brightfield or fluorescence).

***CRITICAL STEP:*** *When sections from the same animal are mounted on different slides (e.g. sections from wells 1 and 7 of a series of 12), we recommend to separate sections (e.g. using “split scene” in Zen) before import. Section can then be reordered and numbered by adding “_N_1; _N_2…” to the file name. Importing those files to QuPath will considerably facilitate section ordering as well as their subsequent analysis*.

**2. Perform cell detection** □ **Timing**: 1h30-3h30 per Qupath project

We provide scripts that facilitate unbiased cell detection while minimizing variability observed between sections. These scripts use thresholding information within QuPath (i.e. median intensity) or call thresholding methods contained within ImageJ (Otsu, Percentile, Mean…).

a. Click “Automate”, select “show script editor” then “File open”. Choose the script “Automatic_threshold_application_(QuPath).groovy” or “Automatic_threshold_application (ImageJ).groovy” to perform adaptive thresholding using QuPath “median” or ImageJ “Mean”. We recommend using ImageJ “Mean” thresholding method (see results) but users can choose another one. To do so, open script “macro-threshold-auto-c-Fos.ijm” and indicate required imageJ thresholding method to use on line 25 (e.g. Otsu, Percentile…). **?TROUBLESHOOTINGS**
b. Run the project by selecting the sections to be included in the analysis. An automatic and optimal thresholding will be defined for each section, allowing optimal cell detection even if differences in staining intensity are observed between sections. **?TROUBLESHOOTINGS**

***CRITICAL STEP:*** *It is important to not pre-selected a section in the project before running the script in order to avoid its exclusion from the analysis. (file -> recent project -> click on last project opened)*

**3. Produce and apply classifier** □ **Timing**: 5 min per section (i.e. ∼50 minutes for an experiment including 5 mice)

a. Create a separate folder containing 2 randomly chosen sections per animal in the experiment (i.e. sections stained at the same time) in order to train the classifier.
b. Run the desired cell detection method (see above)
c. Select the first section. Detected cells appear in red. Fill color can be remove by pressing “F” or clicking on 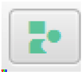
d. Move around the slice (to zoom: use the mouse wheel, to move around the slice: press the spacebar and move by clicking and dragging the mouse) to identify areas with only staining or only artifacts.
e. Select 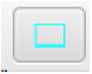 and outline the area where staining or artifacts are located.
f. In the “Annotation tab”, define a class for your outline. To create a class “artefact”, select “present” in the drop-down menu, then click on “set class” to assign it.
g. Repeat the task 5 times for each “class”.
h. At the end, open “Train object classifier” in the “Classify tab”, then “Object Classification”.
i. Click on “Live update” to evaluate the effect of the training on the cell detection.
j. If appropriate, go to the next section or add a new section to upgrade the classifier. We recommend not exceeding 10 sampled areas per class.
k. Once all sections are processed. Go back to the “Train object classifier” tab, in “Load training”, add all processed sections, then click “Apply”.
l. In “Classifier name”, give a name to your trained classifier and save it.

A file is created in the QuPath project folder (folder “classifiers” → “object_classifiers”). This file must be pasted within the “object_classifiers” folder of all animals of the project for automatic quantification.

***CRITICAL STEP:*** *To use classifier within a script, load classifier of interest by selecting it on Load object classifier menu (Classify -> object classification) when one section of your project is opened. Then please follow “workflows” instructions available on QuPath website. (*https://qupath.readthedocs.io/en/stable/docs/scripting/workflows.html#working-with-workflows).

4. **Section registration** □ **Timing**: 2h30 - 3h30 per animal

a. Launch ABBA software :

⍰ Open fiji.exe
⍰ “Plugin” tab -> Biop -> Atlas -> ABBA - ABBA start
⍰ Choose an Atlas (mouse/rat) and slicing mode (coronal/sagittal/horizontal)
⍰ An ABBA window opens
b. Click on “Import” tab then “Import QuPath project” to indicate the location of the QuPath project folder.
c. It is possible to indicate distance between 2 sections at import (if added here, subsequent procedure steps will be easier). All parameters not discussed in this procedure must be applied by default.
d. Following loading, sections appear in the “Slices display” panel on the right.
e. It is possible to show/hide specific section or all of them (with Ctrl-A) and adapt intensit values as follow :

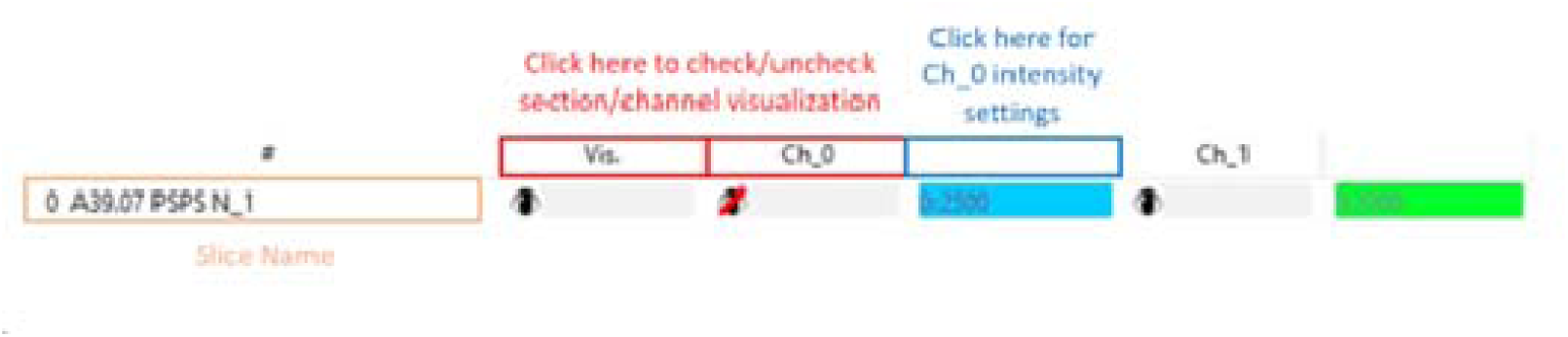

f. On “Atlas display” tab, check Atlas 0 (Nissl) and 1 (Area) then on “Displayed slicing [atlas steps]” distance between sections, if this information has not been added in step “c” (e.g. 24 corresponds to 240µm)

***CRITICAL STEP*** : *Displayed slicing value is an important parameter for optimal atlas registration and will be changed at two differents steps during the procedure :*

⍰ *From step f., displayed slicing value will correspond to the distance between the sections of your series. For example, for 50µm thick sections, if you are sampling every 6th section, the spacing will be 250µm. Thus, a value of 25 will be used. This displayed distance is optimal for initial distribution of brain sections onto the reference Allen Atlas.*
⍰ *Once x/y inclination is done (see step k below), the distance between atlas sections can be reduced so that it corresponds to the thickness of your sections (in our example 5 for 50µm). Such high sampling allows to increase registration precision by refining slices position onto the reference atlas. We recommend choosing a reference section containing well delineated structures such as the Anterior commissure to move all sections (Ctrl-A) at the same time.*
g. Select most rostral and most caudal sections and place them on the corresponding reference atlas sections.
h. Once placed, select these sections and right click to select “define as key slices”. A yellow square appears above them. They are now defined as references for Atlas registration.
i. Select all sections, then use the “Distribute spacing” function found under the “Edit” tab. Small manual adjustments may be necessary to refine the alignment of individual sections with the reference atlas. Indeed, Mistakes can occur during the cutting and collection of serial sections, which may result in slight shifts in section distribution. Although they should be avoided, these shifts should be manually corrected to ensure accurate alignment.
j. To correct small inclinations of the cutting plane, change x (rostrocaudal) and y (left-right) values in the « Atlas Slicing » box (scroll down the window on the right).
k. We recommend using the following sections/landmarks to optimize this step:

- Start adjusting the “Y” axis: for initial adjustments select a section on which the dorsal well defined hippocampi can be seen (i.e. section 70 of the allen brain atlas). To refine, select a second, more rostral section corresponding to the olfactory limb of the anterior commissure (i.e. section 53 of the allen brain atlas). The use of these two structures is particularly sensitive to correct any left-right cutting plane shift. If following this adjustment some sections are identified as being inverted, they can be reverted by clicking the flip symbol within the “Edit Selected Slices” tab.
- Process adjusting the “X” axis: we recommend using a section corresponding to section 80 of the allen brain atlas. In this section, the corpus callosum separates into two parts within each hemisphere. Depending on the rostro-caudal inclination, this separation will be more rostral (with a poorly developed hippocampus laterally) or more caudal (with a very thick corpus callosum)(see supplemental figure 1).
l. As indicated above (“f. Critical Step”), adjust the displayed slicing value to match the thickness of the brain sections (e.g., use 5 for 50µm thick sections). Align all sections as closely as possible to the Allen Atlas. Verify that the distance between sections in the atlas corresponds to the distance between sections in your series. For example, with a 250µm distance, you should observe 5 atlas slices between each of your sections when the displayed slicing value is set to 5. **?TROUBLESHOOTING**
m. Next step corresponds to “Elastix” transformations to refine section registration.

- Select all sections, then click “Elastix registration (Affine)” within the “Align” Tab. We recommend selecting atlas 1 (atlas ARA) and the c-Fos or NeuN channel for optimal results (see **Figure 3B**). Round symbols appear above all sections to indicate the status of the task: yellow signifies that the task is in progress, red indicates that it has not yet been run, and green shows that the task is completed.
- Next, select the “Elastix Registration (Spline)” within the “Align” Tab. Use the same atlas and channel as before. At this stage, the number of reference points should be specified. Note that using a large number of points can be detrimental, potentially causing significant local deformation of the section. We recommend using 10 points.

***CRITICAL STEP*** : *Following section registration is complete, a final check should be performed to ensure that the sections and the reference atlas are correctly aligned. This verification involves two steps:*

- *First, select only Atlas 2 (label border) on the “Atlas display” tab and press “R” to superimpose the Allen atlas onto your brain sections. This view allows experimenters to verify the alignment between the Atlas and the mounted sections and to assess the effectiveness of the Elastix registration steps. If the correspondence Atlas/section is not accurate, the section can be moved rostrally or caudally and the Elastix process can be repeated. (To reset the transformation: right-click on the selected section and choose “Remove last registration” for both spline and affine transformations.*
- *If the overall alignment is good but specific areas (e.g., dentate gyrus) do not match perfectly, the ABBA software allows for local adjustments to improve registration. Use the BigWarp refinement tool by selecting “Edit Last Registration” on the “Align” tab. More details are provided in step n.*

n. BigWarp manual refinement:

Once “Edit Last Registration” is selected, you must specify the atlas (label border (2)) and the channel (c-Fos/NeuN, in our example (1)) to use. Two windows will open: “BigWarp Moving Image” (left) and “BigWarp Fixed Image” (right). Locate the area to be modified (zoom: use the mouse wheel; move around the section: right-click and drag). There are two situations for refinement:

- If a marker (in purple) is already present near the region you want to move, press spacebar to activate “Landmark mode”. Left-click on the marker in the “Fixed Image” window and move it to align the section with the Atlas. **WARNING**: Ensure that landmark mode is off (press the spacebar again) before moving to another region to be adjusted.
- If no maker is present near the region you want to move, you can create a new one. Zoom in on the region to be moved in both windows (after zooming in the first window, press ‘Q’ in the other window to synchronize the fields of view). Place a landmark on the area to be moved in the “moving image” window (press the spacebar and left-click on the area). Then, place a landmark on the corresponding area in the “Fixed Image” window. The region on your section will automatically be adjusted to this new location. Repeat this process as many times as necessary to adapt the sections to the Allen Atlas contours. **?TROUBLESHOOTING**.

***CRITICAL STEP*** *: When using BigWarp, follow those steps for obtaining optimal and reproducible results: 1) realign section borders when necessary, 2) realign interhemispheric “midline”, 3) realign white matter tracts, 4) realign elongated, sinuous brain regions, e.g., dentate gyrus*.

o. Export the atlas registration within QuPath by first selecting all sections, then clicking on “Export registration to QuPath project” located in the “Export” tab). Ensure that the “Erase previous ROI” box is checked before confirming. Yellow circles will appear above the sections and will turn green once the transfer is completed. You can then close the software. **?TROUBLESHOOTING**

**Tips** : Progress through the different steps listed above can be saved at any time by creating an .abba file. We recommend creating this file before exporting atlas registration within Qupath. **WARNING**: please note that once modifications are made to a QuPath project (such as adding or removing sections), it becomes impossible to open a previously saved .abba file. Therefore, exercise caution when making changes to the QuPath project after saving the .abba file.

***CRITICAL STEP:*** *To apply registration on QuPath project, please follow instructions available on ABBA-documentation website: (*https://abba-documentation.readthedocs.io/en/latest/tutorial/4_qupath_analysis.html#automating-the-import-for-all-slices)

5. **Export results on QuPath** □ **Timing**: 1-3 min per animal

a. In “Measure”, select “Export Measurements”.
b. Select sections of interest, then define the save path. Indicate “Annotation” in “Export type” and “comma (csv)” in “Separator”.
c. Add a specific name for the csv file.
d. Click on Export
e. All data for subsequent analyses, are stored in these csv files
6. **Analysis results with R scripts** □ **Timing**: 1-2min

Place all csv files into a single folder and execute the scripts using the following procedure:

a. Qupath_ABBA_analysis_QualityControl.R **?TROUBLESHOOTING**
b. Script for QuPAth_ABBA_analysis_Comparing groups.R (working in cooperation with Atlas creation.R script(**b1**)) **?TROUBLESHOOTING**
c. 3D data representation script. **?TROUBLESHOOTING**

### Timing

The duration of this protocol varies depending on the number of groups, animals, and sections per animal involved in the analysis. For instance, in our example, we consider two groups, each comprising five animals, with approximately 20 sections per animal. The analysis described above was conducted on a Windows 10 machine equipped with 32GB of RAM and QuPath 0.4.3.

Step 1: **QuPath folder creation with czi files included** : 5 min per animal. For reorganization of all czi files and rename before implementing it on the Qupath project.

Step 2: **Cell detection script**: ∼2h for 20 sections

Step 3: **Classifier creation**: around 5 min per section selected. For our analysis we have **2 slices** x **5 animals** X **2 groups** = **20 sections selected** ∼2h

Step 4: **Reference Atlas registration:** Total of 2h30 - 3h30 for one animal

Step 4.f to 4.l Distribute section on reference Atlas (including x/y inclination adaptation) ∼ 1h

Step 4.m Elastix registration ∼ 2-3 min per animal

Step 4.n Manual refinement ∼ 5-8 min per section, around 2h for an entire animal.

Step 5: **Export csv files**: 1-3min per animal; 30min in total in our case.

Step 6: **Script analysis:** 1 to 2 mins

**Total for one animal**: around 7h45.

### Troubleshooting

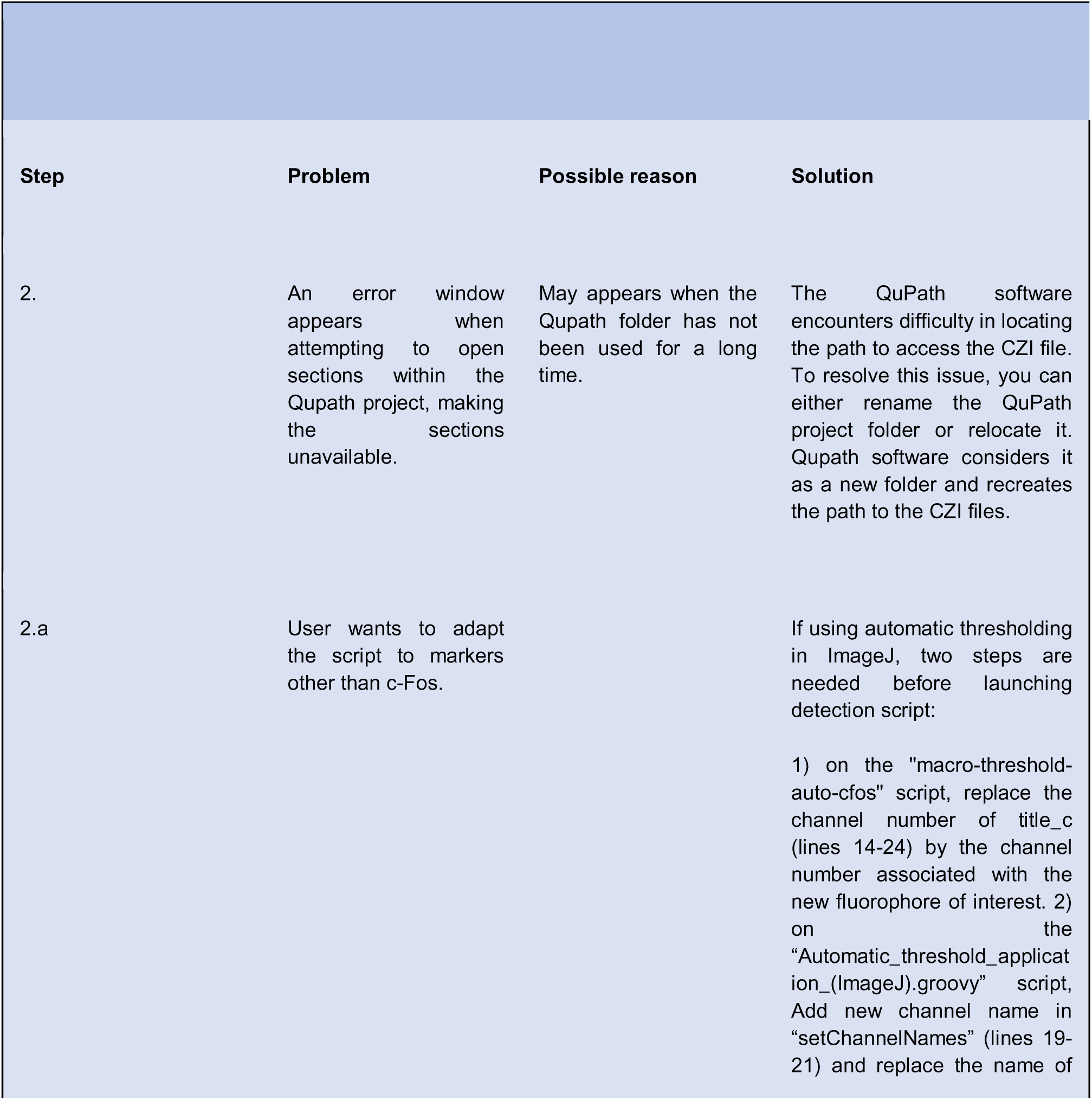
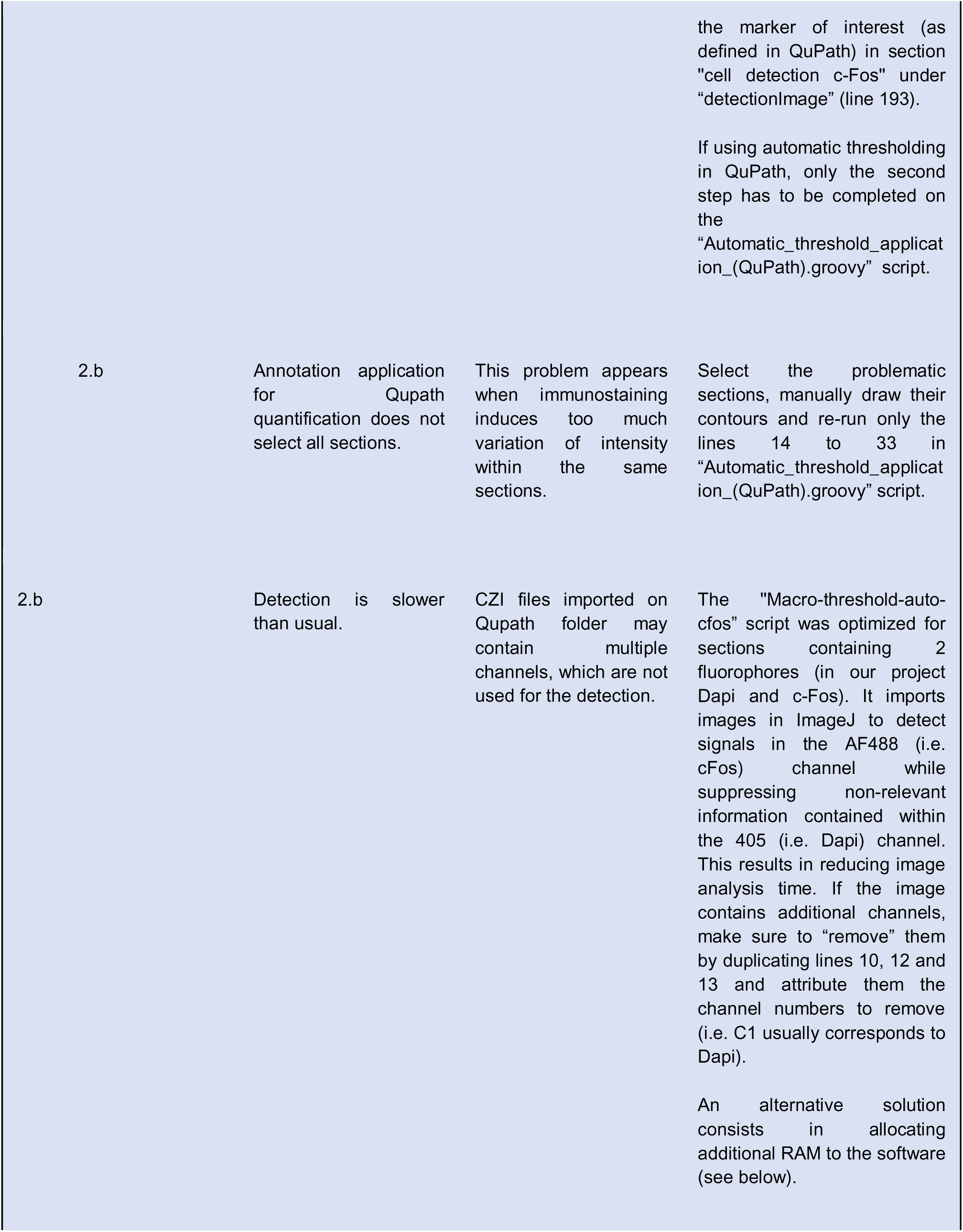
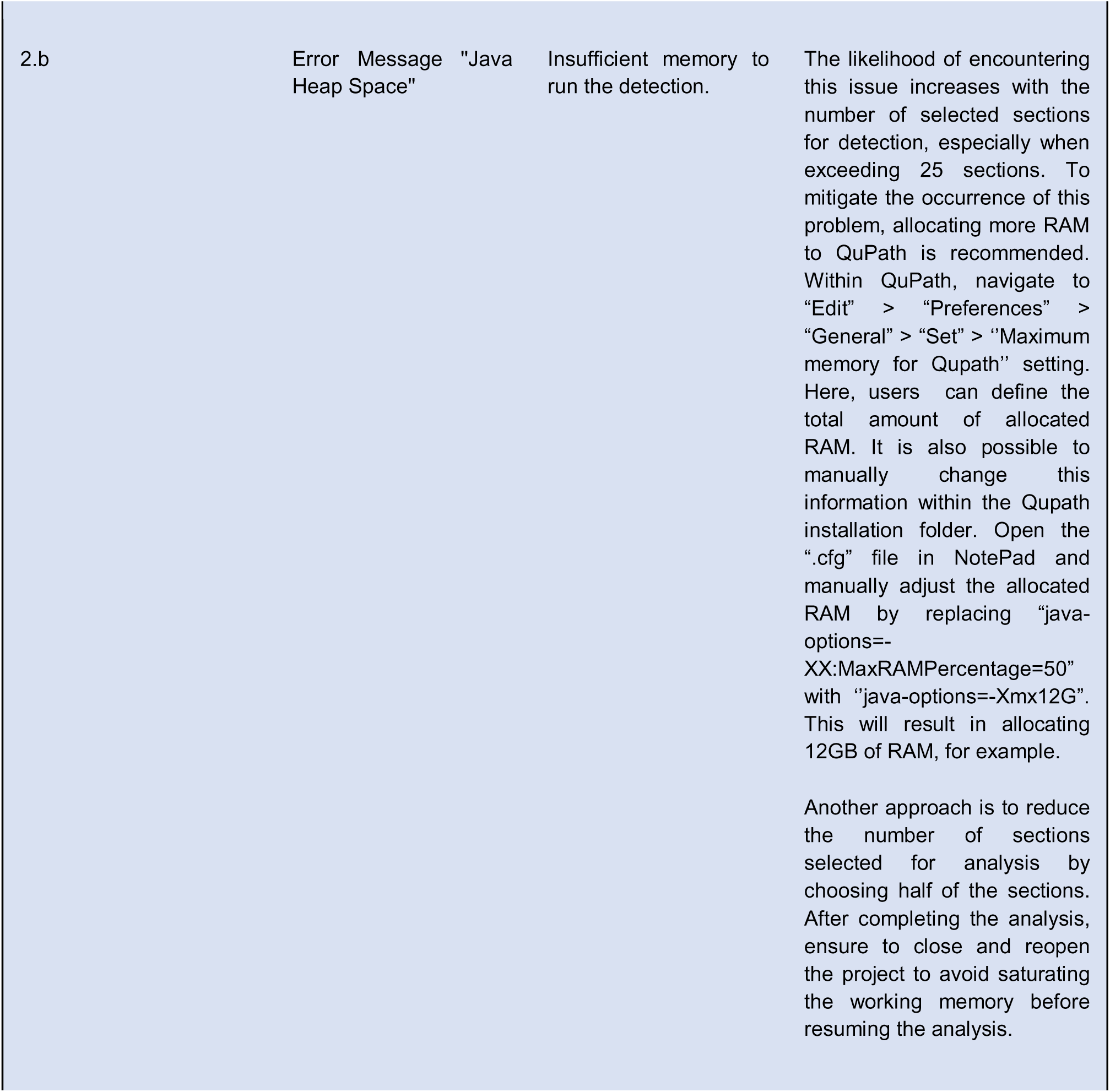
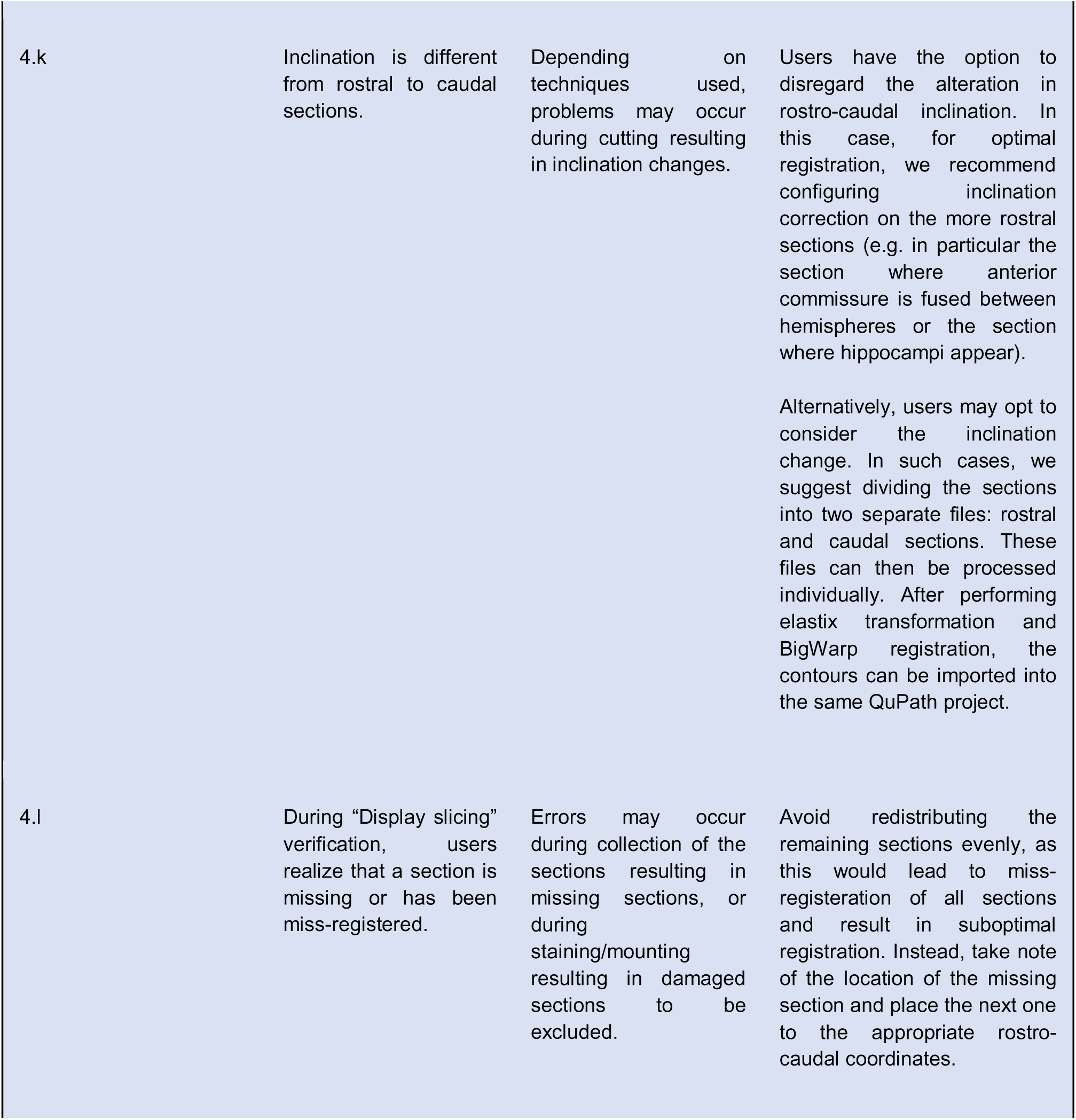
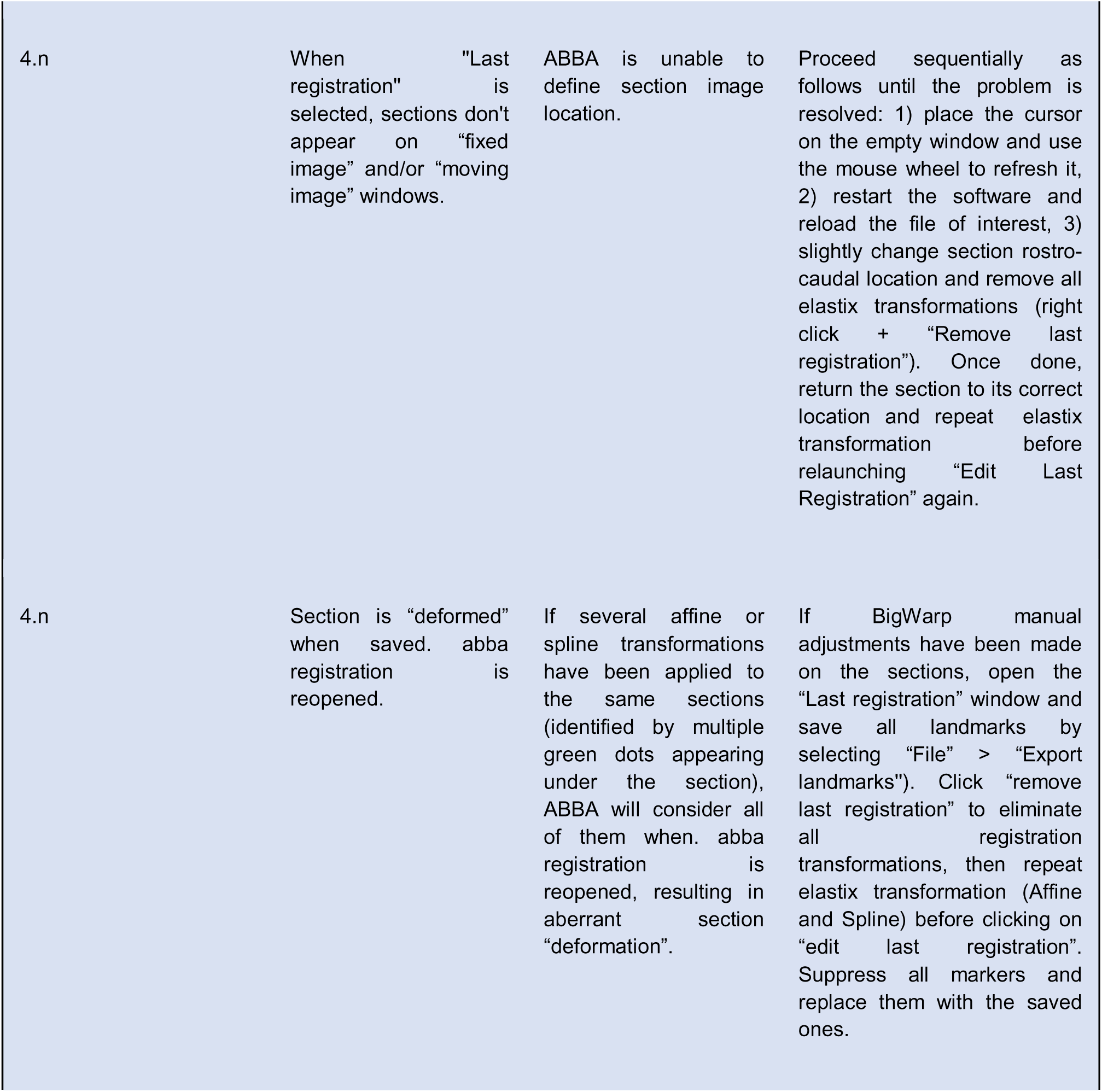
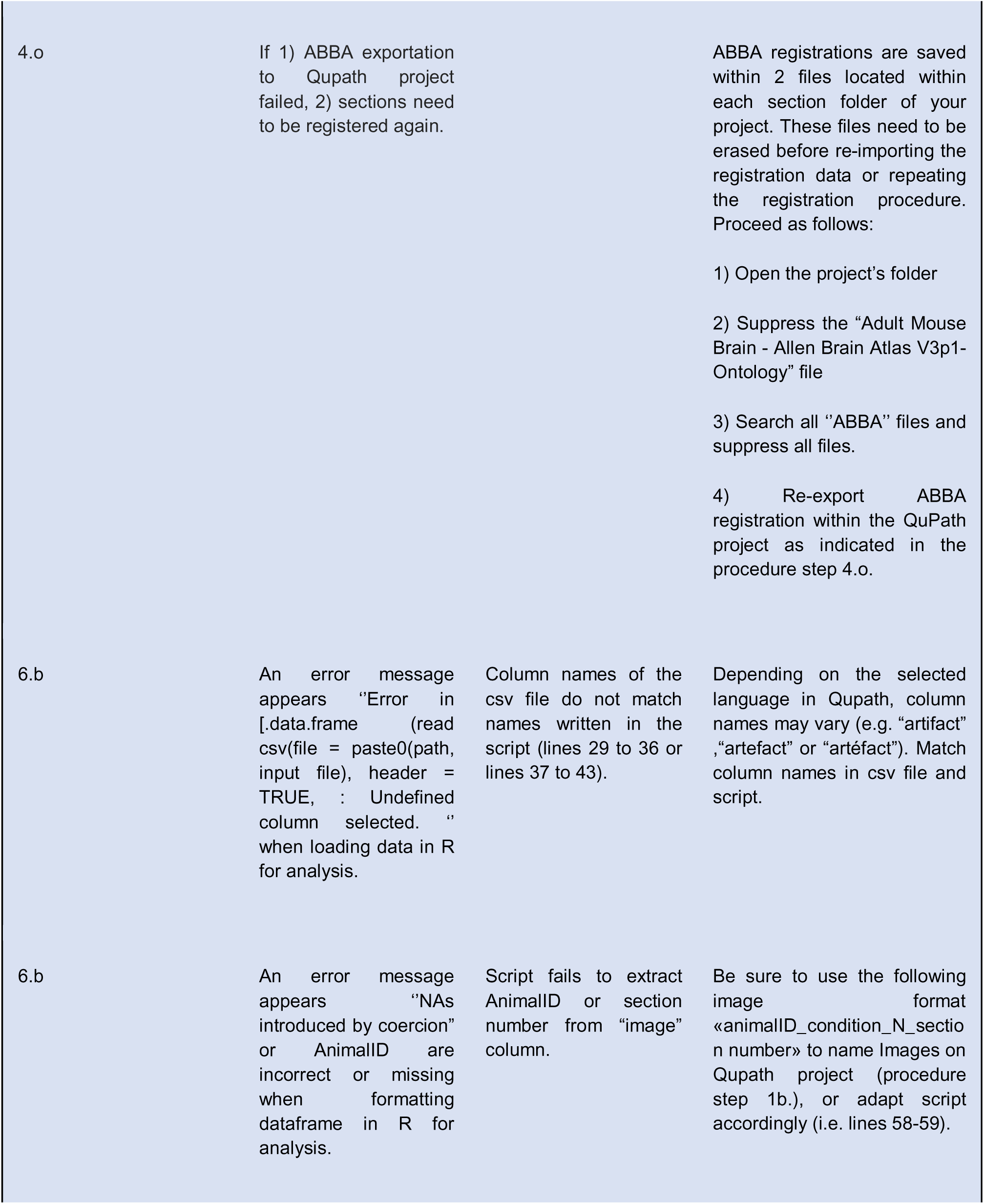
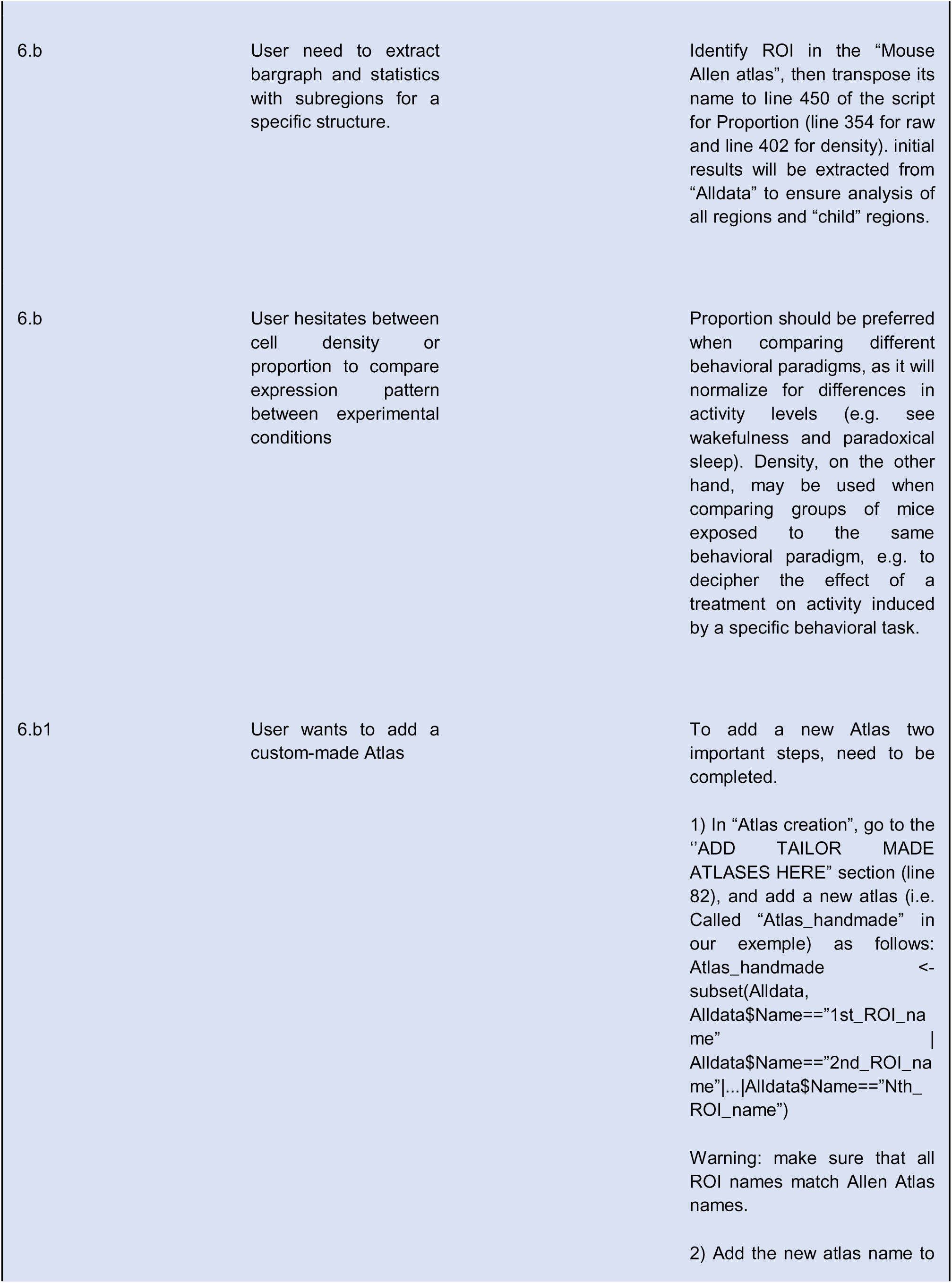
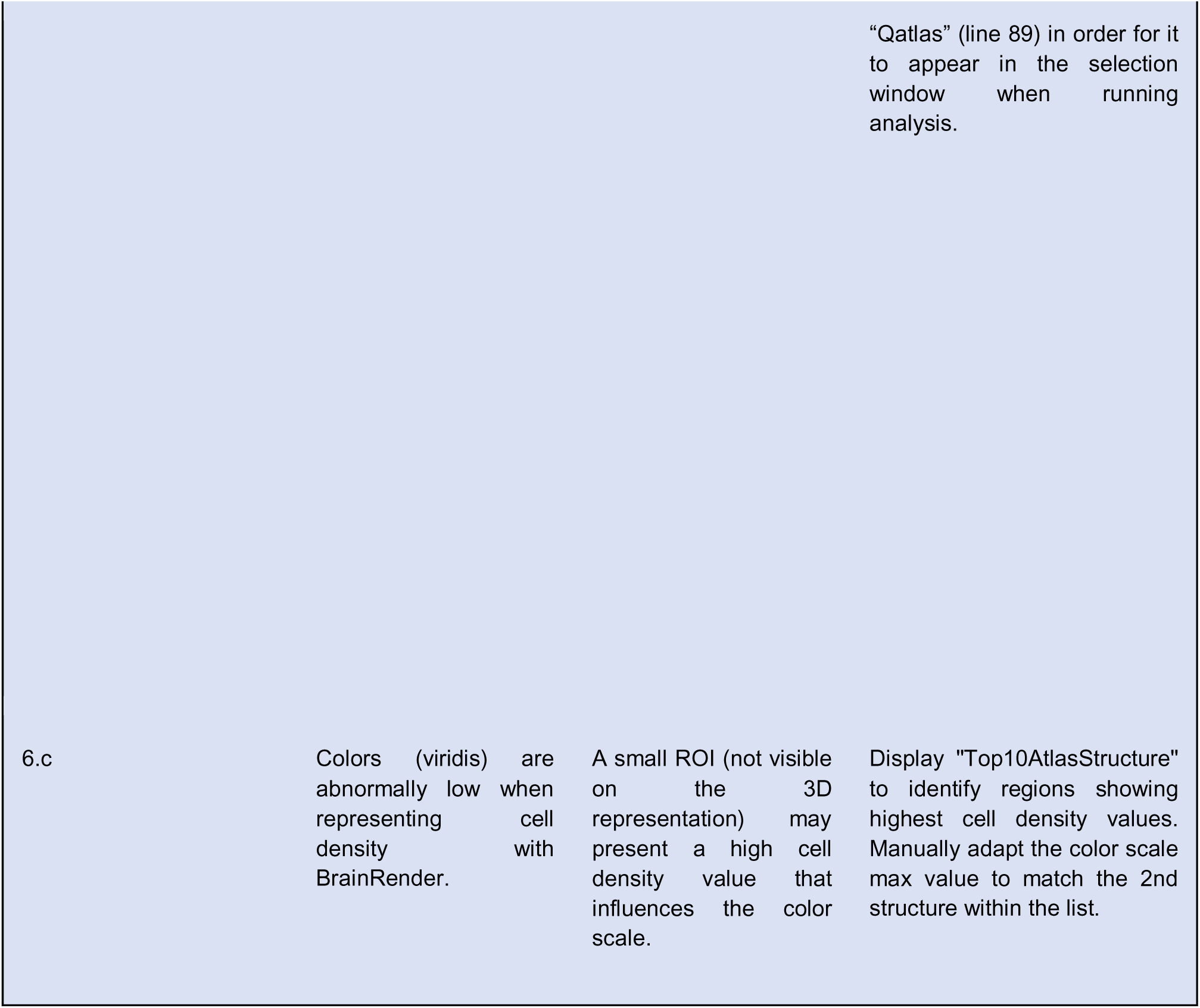

## SUPPLEMENTARY INFORMATION

### -EXTENDED DATA

**Extended Data Figure 1: Reference point for precise cutting plane correction.**

**a)** Refinement of X-axis, corresponding to the dorsoventral axis, using the corpus callosum (CC). The red box highlights the dorsal CC region where modifications are most visible with varying cutting planes, allowing precise adjustment of the atlas representation (bleu-yellow) to match the original section (AF488). The Y-axis, corresponding to the left-right axis, can be refined on the same section by focusing on the hippocampi, whose shapes vary greatly even with minimal shift in the Y-axis. **b)** Y-axis calibration can be further refined on a more rostral section containing the anterior commissure (AC). Yellow lines delineate the AC into three parts, defining landmarks for detecting slight shifts in Y-axis cutting plane.

**Extended Data Figure 2: Quality Control Assessments for Individual Animals.**

Typical quality control outputs for individual animals generated by the script “QuPath_ABBA analysis_QualityControl.R”. The script produces a six-panel figure for each animal CSV file located within the designated folder. **a-b)** Panels detecting missing or aberrant (e.g. folded) sections. **c)** Panel identifying sections with abnormal percentages of artifacts. **d-e)** Panels showing the percentage of c-Fos+ cells within white matter (WM) or ventricles (VS). Values exceeding 5% indicate sections with suboptimal registration. **f)** Panel presenting c-Fos+ cell density within principal brain regions for rapid assessment of abnormal activity patterns between animals exposed to similar experimental paradigms.

**Extended Data Figure 3: 3D Brain Models for qualitative and quantitative representation and comparison of c-Fos expression across experimental conditions.**

**a)** Comma-separated values (.csv) datasheets generated in QuPath are processed with BrainRender to produce 3D brain representations of c-Fos positive cell densities. Results obtained for Paradoxical Sleep (PS) and Wakefulness (WF) are represented using a viridis, yellow (high) to blue (low) scale, using the “Major Division” Atlas. **b)** Direct comparison of results obtained in distinct experimental conditions based on c-Fos positive cell proportions. Results highlight preferential activation in PS (blue gradient) or WF (red gradient) conditions, with regions reaching significance highlighted in yellow (p<0.05) or dark yellow (p<0.001). N=5 mice per group.

**Extended Data Figure 4: Comparison of the three selected thresholding methods on quantifications performed in subcortical regions.**

**a-b**) Comparison of results obtained using the three selected thresholding methods in subcortical structures as represented by bar graphs for c-Fos cells proportion **a**) and 3D brain representations for c-Fos positive cells densities **b**). Note the relative preserved contrast observed between regions when using all thresholding methods, as indicated by values from non-parametric one-way Anova (Krustal-Wallis) statistics showing similar dispersion of c-Fos cells proportions. Abbreviations: DG: dentate gyrus, HIP: hippocampal region, HY: hypothalamus, LHA: lateral hypothalamic nucleus, TH: thalamus, MBO: mamillary body.

**Extended Data Figure 5: Analysis methods for comparing c-Fos activity levels and/or patterns during different behaviors.** a) Quantification of mean number of c-Fos positive cells detected per condition. b-e) Impact of analyses methods for comparing c-Fos activity: b) 3D projections of c-Fos density through different conditions. **c-d)** Bar graphs comparing c-Fos positive cell quantifications using raw cell numbers, density or proportion throughout cortical areas for Wakefulness *vs* Social Behavior (c) and Wakefulness *vs* Paradoxical Sleep (d). **e)** Direct 3D comparison highlighting regions preferentially activated during WF (blue gradient) when compared to regions preferentially activated during SB or PS (red gradient) using the three analysis methods. Data are expressed as Mean ± SEM, *P<0.05; **P<0.01; ***P<0.001 (n=5 mice/condition).

**Extended Data Figure 6: Analysis methods for comparing c-Fos activity levels and/or patterns during different behaviors, second part**. **a-b)** Bar graphs comparing c-Fos positive cells quantifications using raw cell numbers, density or proportion, within CA1, CA2 and CA3 regions of the hippocampus and cortical layers of the ventral retrosplenial (RSPv) for Wakefulness *vs* Social Behavior and Wakefulness *vs* Paradoxical Sleep. **c)** Sagittal and horizontal 3D representations highlighting statistical differences identified throughout the brain in the aforementioned comparisons using the “summary atlas”. Data are expressed as Mean ± SEM, *P<0.05; **P<0.01; ***P<0.001 (n=5 mice/condition).

### Supplementary table

<u>Definition_Allen_atlas_resolutions</u>: This excel file gathers all the structures/sub-structures listed in the Allen Atlas and organizes them according to the three predefined atlases; “Full_Atlas”, “Summary_Structure” and “MajorDivision”. Thus, when MajorDivision is selected for example, only structures checked in MajorDivision column will be selected in the “Script for QuPath ABBA analysis Comparing groups” script.

#### - Source Data

**DATA SOURCE AFIGURE 4:** gathers the data used for figures 4c and 4f to compare automatic thresholding methods, i.e. low (MeanIJ), med (MedQP) or high (MaxE), for condition comparison analyses.

**DATA SOURCE AFIGURE 5:** gathers the data used for condition comparison. Proportion analysis are shown with pval data for every structure listed in “Summary Structures” and “Full List” Atlases.

**DATA SOURCE AFIGURE 6:** gathers the data used for Trap2 tdTomato *vs* c-Fos comparison. Proportion analysis are shown with pval data for every structure listed in “Summary Structures” and “Full List” Atlases.

**DATA SOURCE EXTENDED DATA AFIGURE 5&6:** gathers the data used for condition comparison. Density and raw number analysis are shown with pval data for every structure listed in “Summary Structures” and “Full List” Atlases.

## Author contributions statements

Conceptualization: **OR**, **NM**, **PHL**. Methodology and analysis: **SC** established all histological protocols and performed all analyses of the paper. Writing – original draft: **SC** and **OR**. Writing – review and editing, **SC**, **MB**, **SD**, **PHL**, **NM** and **OR.** Software benchmarking and selection, **MB.** Script editing and refining (Qupath, R-studio, Biorender): **NV**. Workflow testing and refinement: **SC**, **SD**, **NV**. Behavioral testing and tissue production: **SE**, **GMD**, **RM**, **JC**. Funding acquisition, **OR**, **NM**, **PHL**.

## Acknowledgments/Financial support

We thank all zootechnicians from the SBRI and CRNL for their help with mice colonies, as well as the Ciqle Platform for use of equipment and advices. We thank Vincent Rosserro for initial testing of the workflow during his Master. This work was supported by the LABEX CORTEX (ANR-11-LABX-0042) of Université Claude Bernard Lyon 1, within the program “Investissements d’Avenir” (dévision n° 2019-ANR-LABX-02), ANR NeoReGen (ANR-22-CE17- 0029), ANR Rewired (ANR-22-CE16-0019), operated by the French National Research Agency (ANR).

Marine Breuilly is funded by LyMIC and LABEX CORTEX (ANR-11-LABX-0042) of Université Claude Bernard Lyon 1. The LyMIC facility has received funding from the Auvergne Rhône Alpes region and the European Union.

