## Supplementary figures and images for "Establishment of an optimized and automated workflow for whole brain probing of neuronal activity"

### Extended Data Figure 1

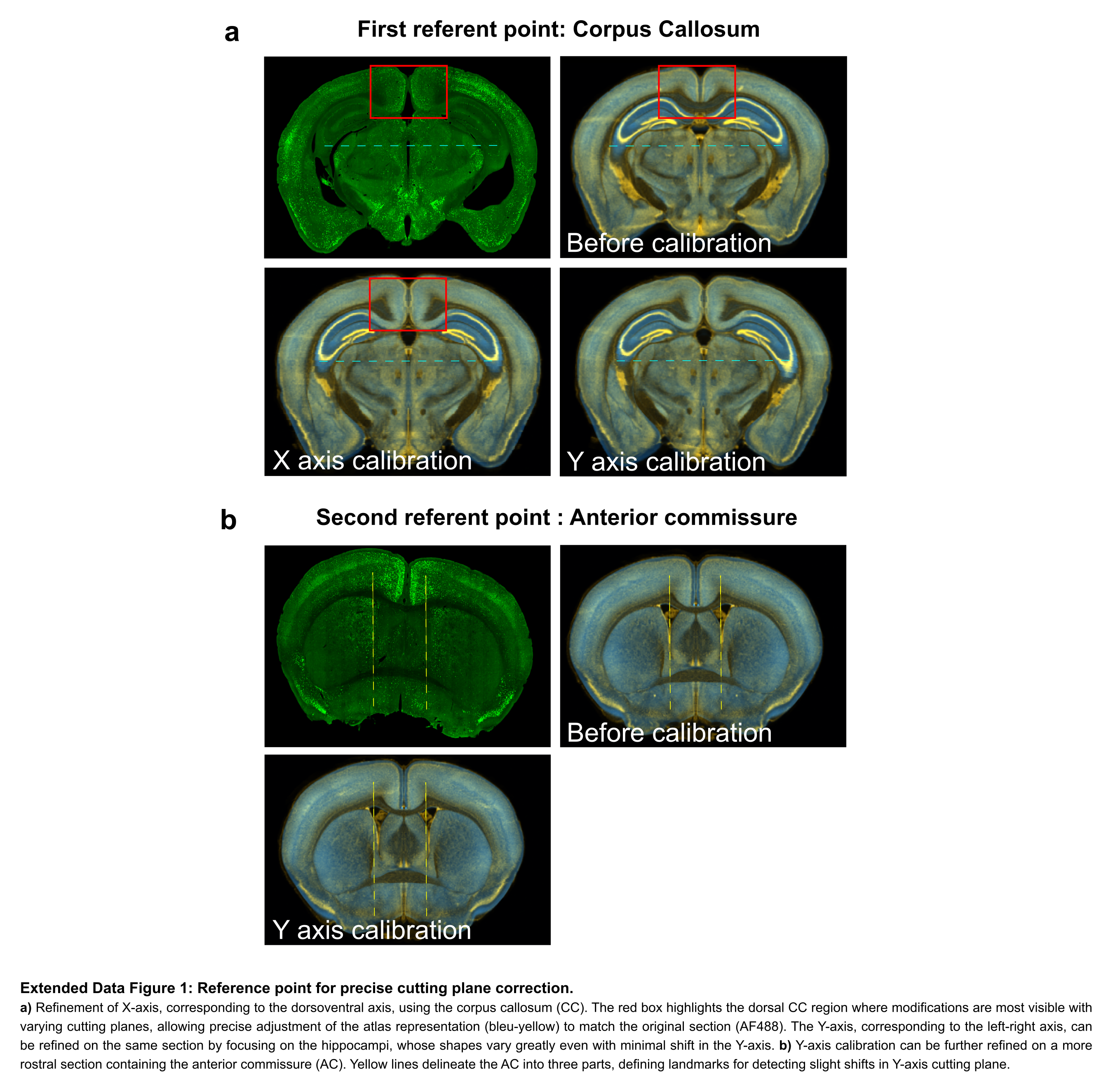

### Extended Data Figure 2

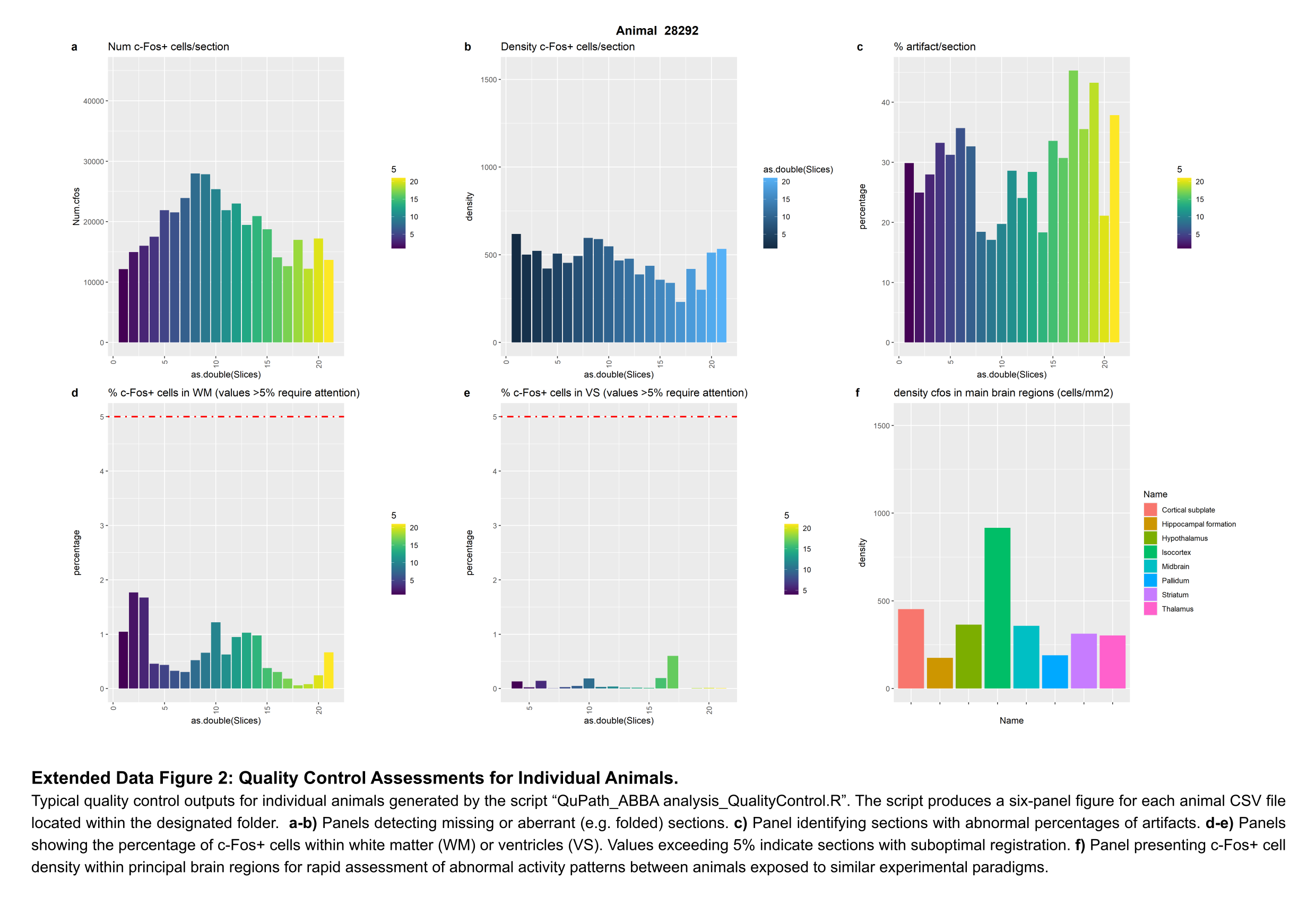

### Extended Data Figure 3

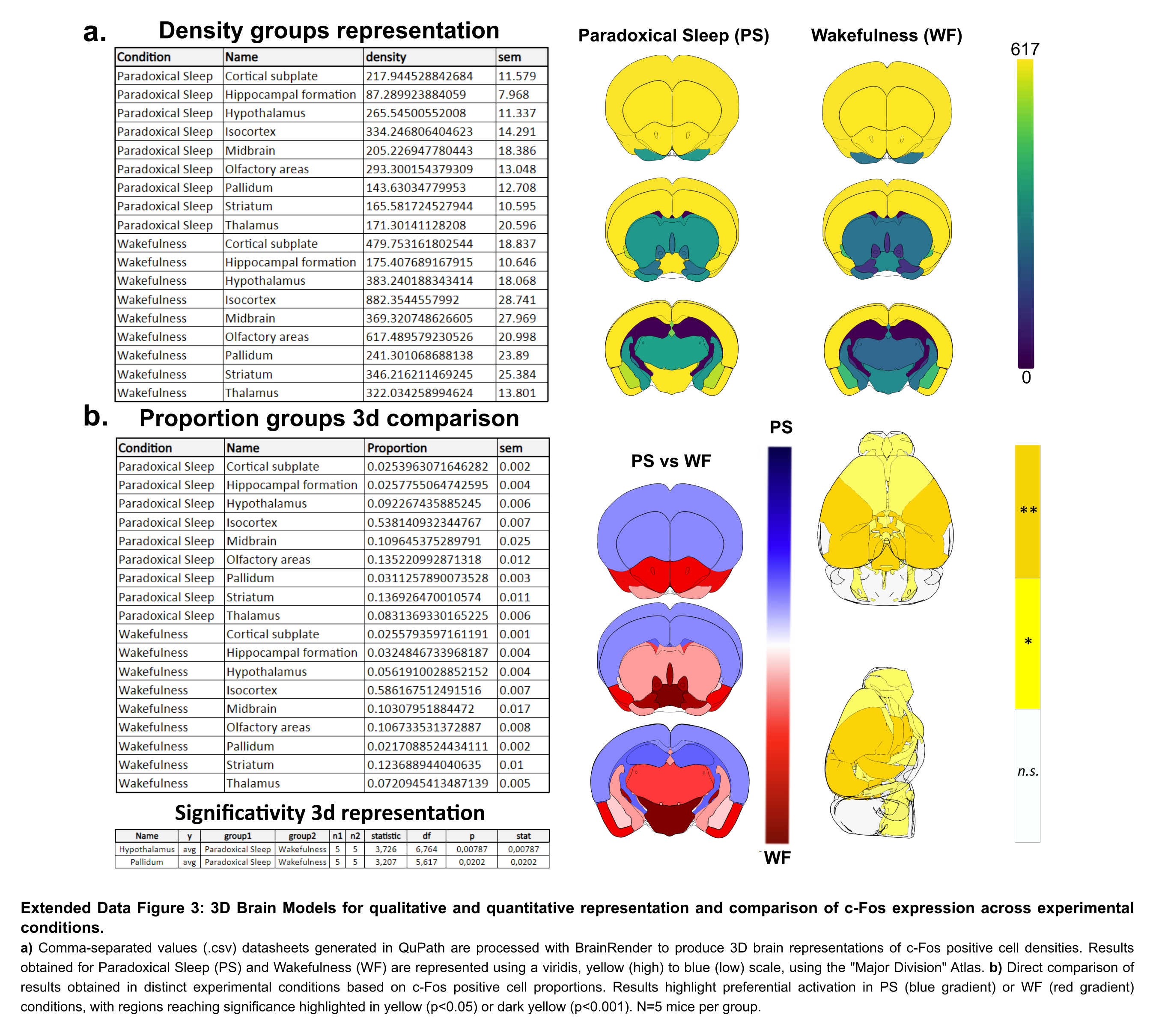

### Extended Data Figure 4

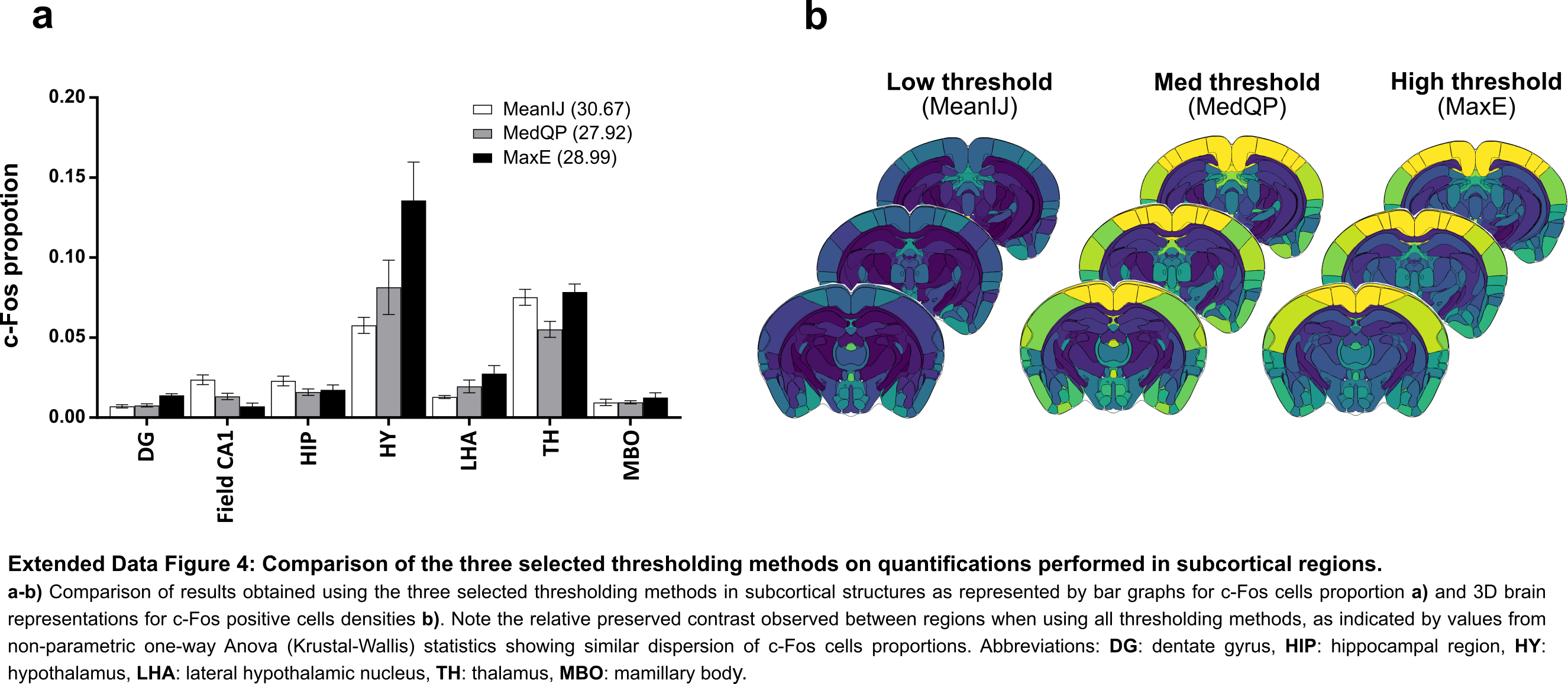

### Extended Data Figure 5

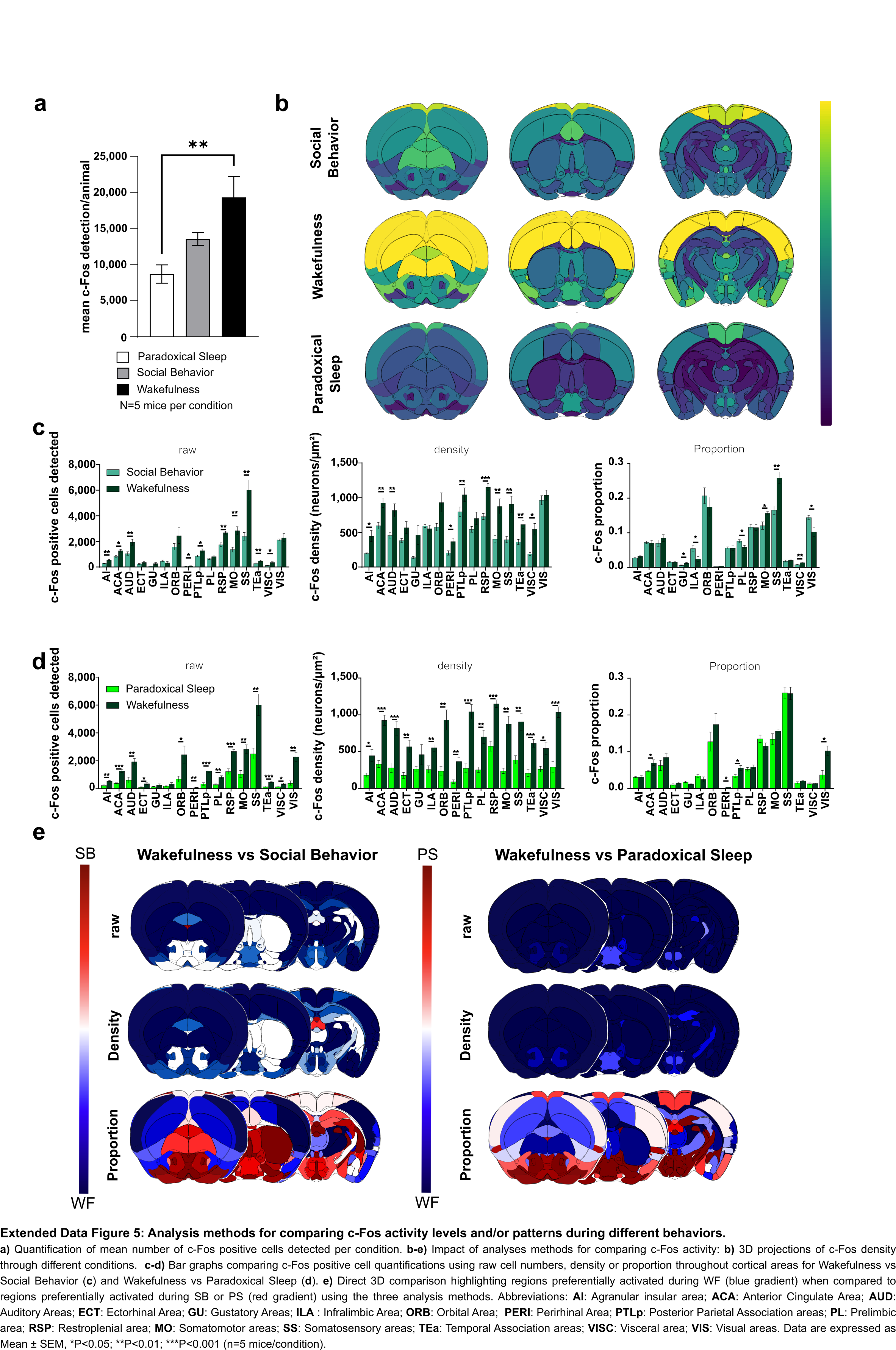

### Extended Data Figure 6

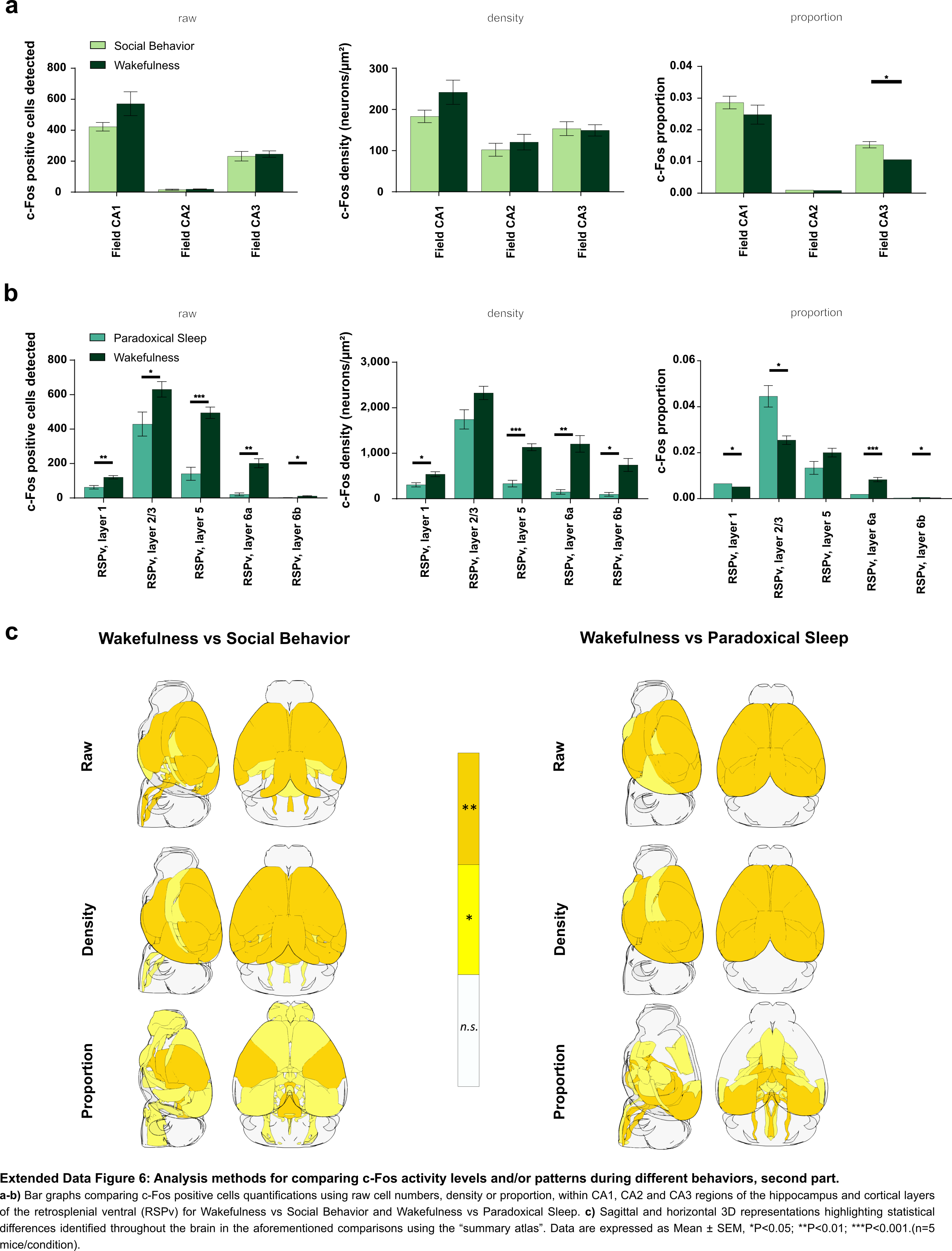
